# Primary cilia mediate diverse kinase inhibitor resistance mechanisms in cancer

**DOI:** 10.1101/180257

**Authors:** Andrew D. Jenks, Simon Vyse, Jocelyn P. Wong, Deborah Keller, Tom Burgoyne, Amelia Shoemark, Maike de la Roche, Athanasios Tsalikis, Martin Michaelis, Jindrich Cinatl, Paul H. Huang, Barbara E. Tanos

## Abstract

Primary cilia are microtubule-based organelles that detect mechanical and chemical stimuli. Although cilia house a number of oncogenic molecules (including Smoothened, KRAS, EGFR, and PDGFR), their precise role in cancer remains unclear. We have interrogated the role of cilia in acquired and *de novo* resistance to a variety of kinase inhibitors, and found that in several examples, resistant cells are distinctly characterized by an increase in the number and/or length of cilia with altered structural features. Changes in cilia length seem to be linked to the lack of recruitment of Kif7 and IFT81 to cilia tips, and result in enhanced hedgehog pathway activation. Notably, Kif7 knockdown is sufficient to confer drug resistance in drug sensitive cells. Conversely, targeting of cilia length or integrity through genetic and pharmacological approaches overcomes kinase inhibitor resistance. The identification of a broad mechanism of pathway-unbiased drug resistance, represents a major advancement in oncology, and helps define a specific and important role for cilia in human cancer.

## Introduction

Primary cilia are microtubule-based sensory organelles that detect mechanical and chemical stimuli, and are formed by nearly all vertebrate cells (Garcia-Gonzalo and Reiter, 2012). These antenna-like organelles house a number of oncogenic molecules including Smoothened, KRAS (Lauth et al., 2010), EGFR, and PDGFR (reviewed in (Christensen et al., 2012)). Although loss of cilia has been associated with the onset of malignancy in some human tumors (reviewed in (Basten and Giles, 2013)), in others, cilia appear to be necessary for cancer cell survival (Han et al., 2009),(Wong et al., 2009),(Li et al., 2016). In fact, depending on the nature of the driver oncogenic lesion, cilia can have opposing roles in tumorigenesis even in the same tumor type. For example, a study by Han *et* al (Han et al., 2009) showed that while removal of cilia prevented tumor growth in a mouse model of medulloblastoma (MB) driven by transgenic expression of a constitutively active form of Smoothened (Smo), the same perturbation in a MB model driven by expression of constitutively active GLI2 promoted tumor growth. Therefore, the role of cilia in cancer remains unclear, and is likely to be context dependent. Furthermore, characterization of primary cilia in glioblastoma cells suggest that cancer-associated cilia may be structurally distinct (Moser et al., 2014).

A number of oncology drugs target cilia-resident proteins such as EGFR and PDGFR (Christensen et al., 2012). These drugs (e.g. the EGFR inhibitor erlotinib) promote significant tumor regressions in appropriate patient populations (e.g. EGFR-mutant non-small cell lung carcinoma patients). However, these responses are invariably followed by the emergence of lethal drug-resistant disease. Our understanding of the molecular mechanisms of drug resistance has facilitated the design and deployment of second line therapies that can target drug-resistant tumors (Awada et al., 2015). However, the characterization of these mechanisms (particularly those that do not involve mutation of the drug target itself) has been limited, and specific to the individual target or drug.

Thus, drug resistance remains the main obstacle in delivering long-lasting therapeutic benefit. The identification of cell biological processes that facilitate and support the emergence of drug resistance may provide new therapeutic opportunities with broad applicability.

In this study, we report that the number and length of primary cilia are upregulated both in *de novo* and acquired kinase inhibitor resistance. These changes are associated with distinct molecular features at the cilium, including 1) failure to recruit Kif7 and control cilia length, 2) increased Hedgehog pathway activation, and 3) cilia fragmentation. Cilia elongation via Kif7 knockdown is sufficient to increase survival in the presence of kinase inhibitors, thus suggesting that cilia elongation has a critical role in promoting drug resistance. Notably, targeting ciliary pathways or impairing ciliogenesis through downregulation of ciliary proteins can overcome resistance in all cases studied. Thus, we have uncovered a previously unrecognized role for cilia in cancer that provides a rationale for targeting ciliogenesis as a broadly applicable strategy to overcome drug resistance.

## Results

The role of primary cilia in human cancer is ill-defined. Given the wide range of oncogenic proteins that are regulated by or localized to cilia (Christensen et al., 2012) (Lauth et al., 2010), we hypothesized that changes in ciliogenesis could play a permissive role in the emergence of drug resistance. First, we examined EGFR-inhibitor resistance in the EGFR-mutant non-small cell lung carcinoma (NSCLC) cell line HCC4006. We chose this model system because EGFR inhibitors are effective in the treatment of EGFR-mutant lung cancer patients, but resistance to these drugs is inevitable. Furthermore, the mechanisms of drug resistance are still unknown for a large number of these patients. We examined ciliogenesis in these cells by staining for acetylated tubulin, a marker for cilia, or Arl13B, a marker specific for ciliary membranes (Caspary et al., 2007) (Cevik et al., 2010). Interestingly, whereas control HCC4006 cells completely lacked primary cilia, erlotinib-resistant HCC4006 cells generated by chronic exposure to erlotinib – ((Saafan et al., 2016) and Supplementary Fig.1A) showed robust staining for ciliary markers (Fig. 1A).

**Figure 1.**
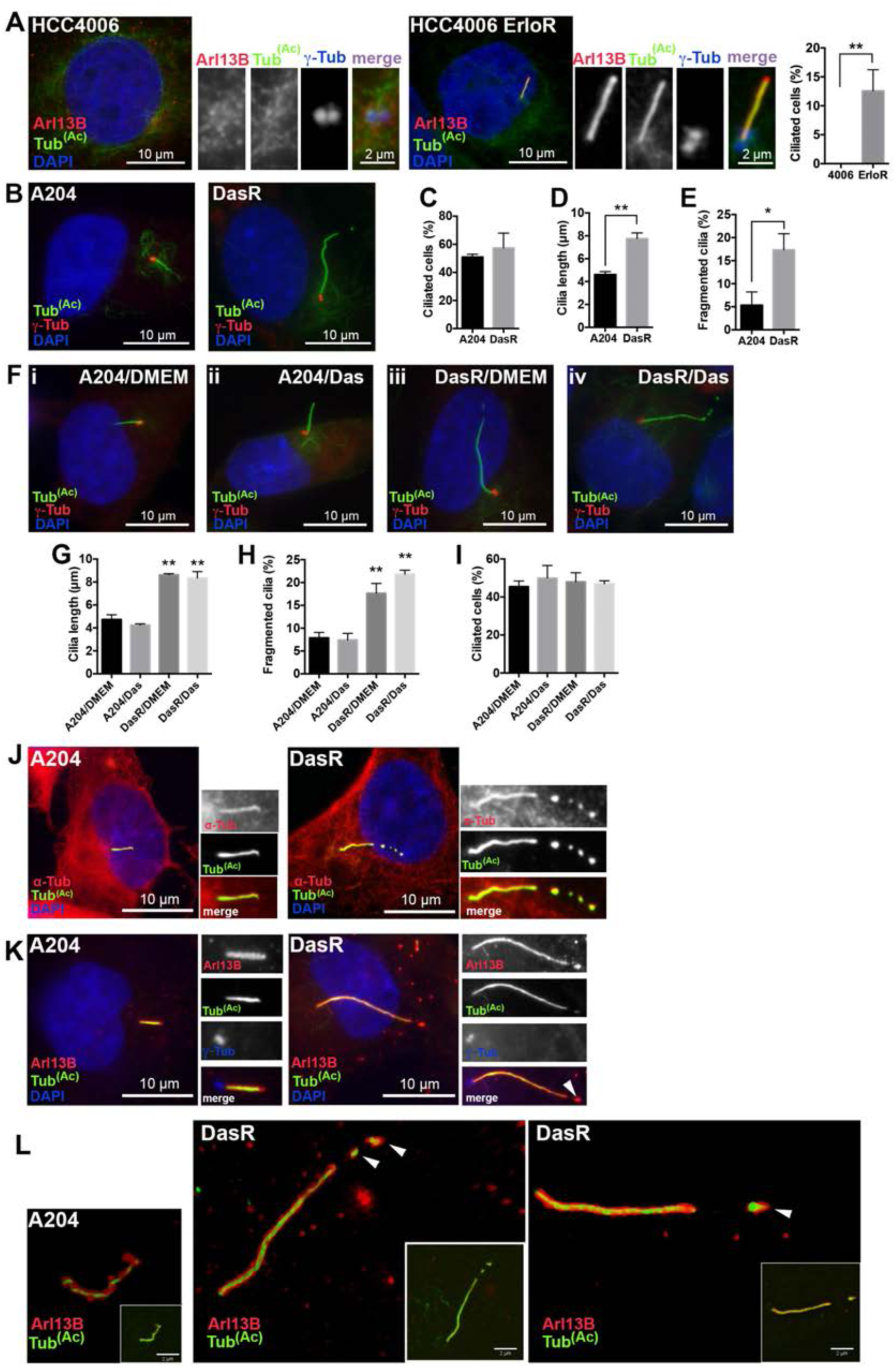
Acquired resistance to kinase inhibitors in patient-derived tumor cell lines shows increased cilia frequency, cilia length and cilia-tip fragmentation. (**A**) Control (left panels) or erlotinib resistant (ErloR) (right panels) HCC4006 lung-adenocarcinoma cells were serum starved for 48 hours to induce ciliogenesis, then fixed and stained with antibodies for acetylated tubulin (green) and Arl13B (red) to mark cilia, γ-tubulin (blue/inset) for centrioles and 4, 6-diamidino-2-phenylindole (DAPI) (blue) to mark DNA. Note that primary cilia were absent from HCC4006 cells but surprisingly are present in the erlotinib-resistant subline. Quantification of ciliated cells is shown on the right. n = 300. Error bars represent s.d. p<0.005, unpaired T test. (**B**) Rhabdoid tumor A204 cells (left panel), or a dasatinib resistant (DasR) subline (right panel) were stained with acetylated tubulin to mark cilia (green), γ-tubulin (red) and with DAPI (blue). (**C**) Cilia quantification for the experiment shown in **B** (n = 300). (**D**, **E**) Cilia length (n = 150) and cilia fragmentation (n = 150) for the experiment shown in **B**. Note that DasR cells show increased cilia length and cilia fragmentation. Error bars represent the s.d. p<0.0007 for **D** and p<0.011 for **E,** unpaired T test. (**F**) A204 or DasR cells were grown in DMEM (i, iii) or DMEM+dasatinib (ii, iv) for 48 hours, then serum starved with (ii, iv) or without (i, iii) dasatinib for 48 hours and stained with acetylated tubulin (green),γ-tubulin (red) and with DAPI (blue). (**G**) Quantification of primary cilia length, (**H**) cilia fragmentation and (**I**) percentage of ciliated cells shown in **F**. n = 150 cilia. The error bars represent the s.d. p<0.0001 for G and H, Tukey’s multiple comparison test, statistical significance calculated by comparing DasR/DMEM and DasR/Das to A204/DMEM and A204/Das. (**J**) A204 (left) or DasR (right) cells were serum starved to induce ciliogenesis, then fixed and stained forα-tubulin (red) to mark all microtubules, acetylated tubulin (green) for cilia and DAPI for DNA (blue). Note thatα-tubulin is present along the entire cilium axoneme in both A204 and DasR cells and it follows cilia fragmentation in DasR cells (right). (**K**) A204 (left) or DasR cells (right) were stained with Arl13B for ciliary membranes (red), acetylated tubulin (green), γ-tubulin (blue/inset) and DAPI (blue). Arrow indicates cilia fragments marked by both acetylated tubulin and Arl13B in DasR cells. (**L**) 3D structured illumination images of A204 and DasR cilia; Arl13B is shown in red and acetylated tubulin in green. Note that at this resolution, Arl13B signal surrounds acetylated tubulin. Arrows indicate budding fragments in DasR cells that contain membrane around them, suggesting an active budding event. These data are representative of three independent experiments.

**Figure 2.**
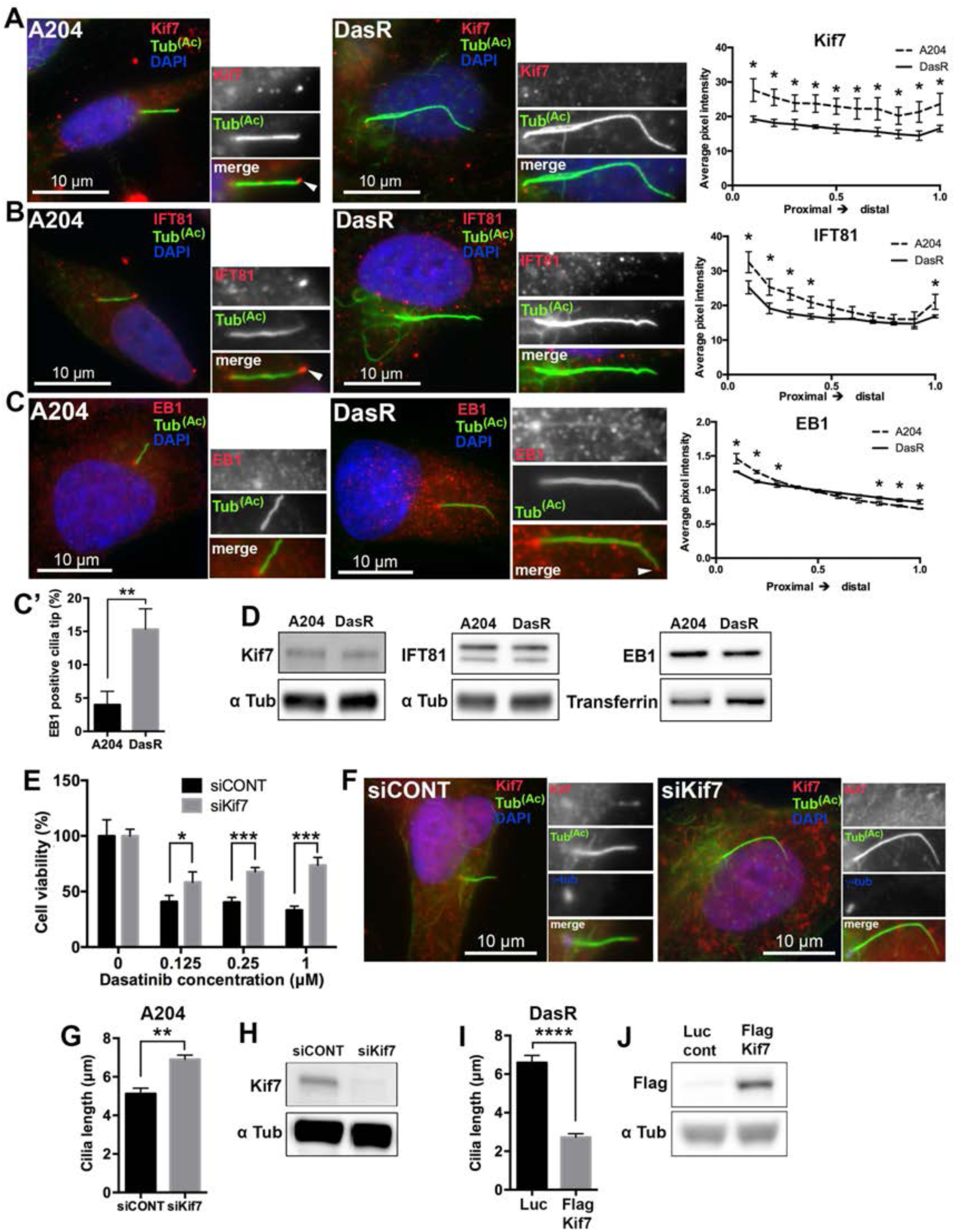
Cilia length control is critical for the acquisition of resistance. (**A**) A204 (left) or DasR cells (right) were serum starved for 48 hours to induce ciliogenesis. After fixation, cells were stained with antibodies against acetylated tubulin (green), KIF7 (red) and with DAPI (DNA, blue). (**B**) A204 (left) and DasR cells (right) treated as in **A**, were stained with antibodies for acetylated tubulin (green), IFT81 (red) and with DAPI (blue). Note that both Kif7 and IFT81 are present along the cilia and at the cilia tip in control A204 cells (arrows) but are absent in DasR cells. Quantification of Kif7 (**A**) and IFT81 (**B**) is shown on the right. n = 150. Error bars represent s.d. Kif7 P-values, proximal to distal: <0.02 <0,01, <0.02, <0.03, <0.03, <0.03, <0.03, <0.03, <0.03, <0.02 (**A**), IFT81 P-values proximal to distal: <0.03, <0.02, <0.001, <0.02, <0.04 (**B**), for an unpaired T test. (**C**) A204 cells (left panel), or DasR (right panels) were stained with acetylated tubulin to mark cilia (green), EB1 (red) and with DAPI (blue). Note that EB1 is present at the cilia tip of DasR cells (arrow/inset) but not in parental A204 cells. Quantification of EB1 shown on the right (Fluorescence intensity is presented as a ratio of total cilia fluorescence intensity). n = 150. Error bars represent s.d. EB1 P-values, proximal to distal: <0.02, <0.002, <0.03, <0.02, <0.005, <0.006 (**C**) for an unpaired T test. (**C’**) EB1 was visually confirmed at the cilia tip of A204 and DasR cells. Chart shows quantification. n=150. Error bars represent s.d. p<0.006, for an unpaired T test. (**D**) Western blots showing total protein levels of Kif7, IFT81 and EB1 (upper panels, indicated) and loading controls (lower panels) in A204 and DasR cells. (**E**) Cell viability (Cell titer Glo) of A204 cells grown in normal media (with the addition of DMSO) or dasatinib (indicated), transfected with either control siRNA or Kif7 siRNA (indicated). Cell viability was normalized to DMSO control treated cells (n = 4); error bars represent s.d. p<0.02 (0.125 μM), p<0.0001 (0.25 μM), p<0.0001 (1 μM), unpaired T test. (**F**) A204 cells were serum starved for 48 hours to induce ciliogenesis, then fixed and stained with antibodies for acetylated tubulin (green) and Kif7 (Boehlke et al.), and with DAPI (blue). Note that A204 cells transfected with a Kif7 siRNA (siKif7) had increased cilia length compared to cells treated with a control siRNA (siCONT). (**G**) Quantification of cilia length shown in **F**. Cilia length in A204 cells transfected with siKif7 was significantly increased compared to siCONT. n = 150, error bars represent s.d. p<0.002, for an unpaired T test. (**H**) Western blot showing Kif7 expression in A204 cells transfected with control siRNA or Kif7 siRNA (indicated) for the experiments shown in **E, F, G**. (**I**) Cilia length quantification of DasR cells expressing Kif7. Expression of FLAG-tagged Kif7 reduced cilia length compared to a non-targeting luciferase (Luc) control plasmid. n = 90, error bars represent s.d. p<0.0001, for an unpaired T test. (**J**) Western blot showing FLAG-Kif7 expression in DasR cells transfected with control luciferase plasmid or FLAG-Kif7 (indicated). These data are representative of three independent experiments.

We then asked whether changes in ciliogenesis could be seen in additional models of drug resistance, where the primary target was not EGFR. To do this, we examined the Rhabdoid tumor cell line A204, which is exquisitely sensitive to the tyrosine kinase inhibitor dasatinib, and a dasatinib-resistant (DasR) subline which we recently characterized and was generated through chronic drug exposure ((Wong et al., 2016) and Supplementary Fig. 1E). Notably, we found that compared to parental cells, DasR cells showed increased cilia length and tip fragmentation (Fig. 1B-E). These effects were neither acute nor transient, as short term treatment with dasatinib did not promote these changes, and withdrawing dasatinib from DasR cells for several days did not revert the effect (Fig. 1F-I).

We also examined ciliation in the EML4-ALK-fusion-positive lung cancer cell line H2228, which is highly sensitive to the ALK inhibitor NVP-TAE684, and a drug-resistant derivative generated through chronic NVP-TAE684 exposure. Similar to HCC4006 and A204, drug resistant H2228 cells showed increased cilia length and number (Figure 3E-G). Of the 5 isogenic models of acquired drug resistance we interrogated, only one (PC-9 lung adenocarcinoma cells) did not show alterations in cilia. In this model, however, cells with acquired resistance to the irreversible EGFR inhibitor afatinib showed a nearly complete biochemical insensitivity to the drug, consistent with the presence of a drug-binding-interfering mutation, a known and common mechanism of drug resistance (Wu et al., 2016). PC9 cells that were resistant to erlotinib exhibited drug-resistant MAPK activity, which has also been described as a mechanism of acquired EGFR inhibitor resistance in both PC9 cells and in lung cancer patients (de Bruin et al., 2014). Therefore, our models seem to cover a range of drug resistance mechanisms. Table 1 summarizes cilia changes identified in all models examined.

**Table 1:** Cilia and drug resistance. Summary of cilia changes observed in resistant cell lines.

| Cell lines/Acquired resistance | Drug resistance | Cilia length | Ciliated cells |
| --- | --- | --- | --- |
| A204 DasR1 | Dasatinib | ++ | - |
| A204 DasR2 | Dasatinib | ++ | - |
| H2228 | NVP-TAE | ++ | ++ |
| HCC4006 | Erlotinib | ++ | ++ |
| PC9 Afatinib | Afatinib | - | - |
| PC9 Erlotinib | Erlotinib | - | - |

| Cell lines/*de novo* resistance | Drug resistance | Cilia length | Ciliated cells |
| --- | --- | --- | --- |
| A549 | Trametinib | ++ | ++ |
| H23 | Trametinib | ++ | ++ |
| H1792 | Trametinib | - | ++ |

| Cell lines/Chemical resistance (acquired) | Drug resistance | Cilia length | Ciliated cells |
| --- | --- | --- | --- |
| A549 | Cisplatin | - | ++ |
| A549 | Carboplatin | - | - |
| A549 | Vinflunine | ++ | ++ |

**Figure 3.**
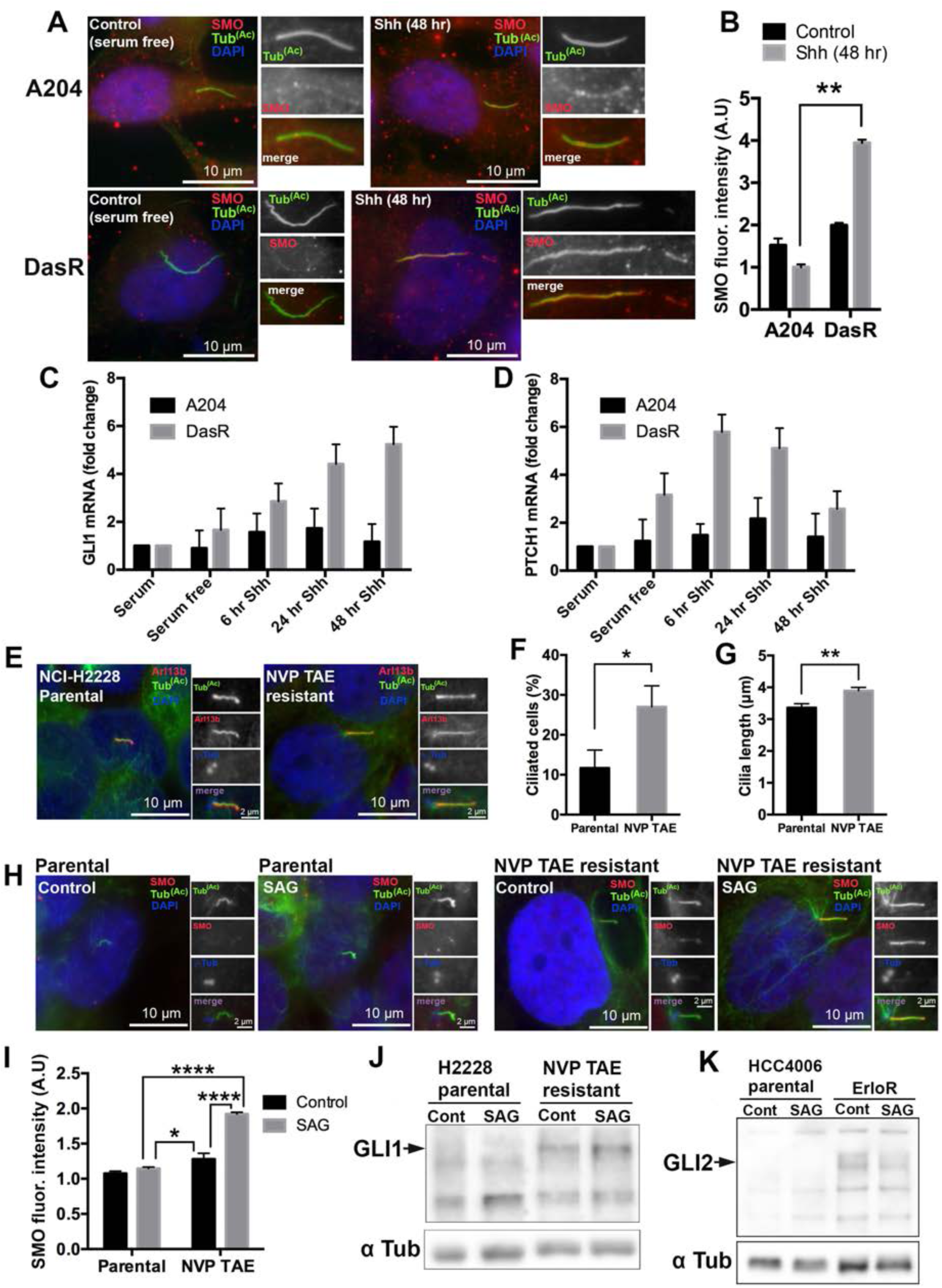
Kinase inhibitor resistant cells show increased Hedgehog pathway activation. (**A**) A204 cells (top panels), or a dasatinib resistant subline (DasR) (lower panels) were serum starved for 24 hours, then either left in serum-free media (left) or treated with human sonic hedgehog (Shh, 5 μg/ml) (right) for an additional 48 hours. Cells were then fixed and stained for acetylated tubulin to mark cilia (green), smoothened (SMO, red) and with DAPI (blue). DasR cells show increased SMO localization to cilia compared to control A204 cells. (**B**) Quantification of SMO cilia fluorescence intensities for the experiment shown in **A**. Fluorescence intensity was normalized to surrounding fluorescence. n = 150, error bars represent s.d. p<0.01, for an unpaired T test. This is representative of three independent experiments. (**C**, **D**) Quantitative polymerase chain reaction (qPCR) showing mRNA levels of Hh target genes of *GLI1* (**C**) and *PTCH1* (**D**) in A204 and DasR cells before serum starvation, after 24hr of serum starvation and after stimulation with 5 μg/ml Shh for the times indicated. TATA box–binding protein (*TBP*) was used as a reference gene and fold change was calculated by comparing mRNA levels relative to control (serum). (**E**) Parental (left panels) or NVP TAE resistant (right panels) NCI-H2228 lung-adenocarcinoma cells were serum starved for 48 hours to induce ciliogenesis, then fixed and stained with antibodies for acetylated tubulin (green) and Arl13B (red) to mark cilia, γ-tubulin (blue/inset) for centrioles and 4, 6-diamidino-2-phenylindole (DAPI) (blue) to mark DNA. Note that primary cilia were shorter in parental cells but longer in the NVP TAE-resistant subline. (**F, G**) Quantification of ciliated cells (F) and cilia length (G) shown in **E**. n = 300 for **F** and n = 150 for **G**. Error bars represent s.d. p<0.02 (**F**) and p<0.005 (**G**), for an unpaired T test. (**H**) NCI-H2228 parental or a NVP TAE resistant subline were serum starved for 24 hours, then either left in serum-free media or treated with human SAG (100 nM) for an additional 48 hours. Cells were then fixed and stained for acetylated tubulin to mark cilia (green), smoothened (SMO, red) and with DAPI (blue). NVP TAE resistant cells show increased SMO localization to cilia compared to parental cells. (**I**) Quantification of SMO cilia fluorescence intensities for the experiment shown in **H**. Fluorescence intensity was normalized to surrounding fluorescence. n = 150, error bars represent s.d. p<0.0001 (parental SAG vs NVP TAE SAG), p<0.004 (parental SAG vs NVP TAE control) and p<0.0001 (NVP TAE control vs NVP TAE SAG), for Tukey’s multiple comparison test. (**J**) Western blot showing GLI1 expression in H2228 parental cells and the NVP TAE resistant subline. Cells were serum starved for 24 hours, then either left in serum-free media or treated with human SAG (100 nM) for an additional 48 hours. Note the increased expression of Gli1 in NVP TAE resistant cells compared to parental cells. (**K**) Western blot showing Gli2 expression in HCC4006 parental cells and the ErloR resistant subline. Cells were serum starved for 24 hours, then either left in serum-free media or treated with human SAG (100 nM) for an additional 48 hours. Note the increased expression of Gli2 in ErloR resistant cells compared to parental cells. Data shown are means ± SD, n = 3 independent experiments.

Cilia-derived vesicles have been shown to have important intercellular functions in tetrahymena and Chlamydomonas (Wang and Barr, 2016; Wood et al., 2013). However, cilia-derived fragments have never been described in cancer cells.

We therefore set out to characterize the nature of the observed cilia fragmentation in drug-resistant cells by examining tubulin post-translational modifications. No difference was observed in total tubulin acetylation, or detyrosination. However, the extent of tubulin polyglutamylation along the cilia was reduced in DasR cells (Supplementary Fig. 2A-C). Furthermore, staining acetylated tubulin together with alpha-tubulin, to mark all microtubules, clearly showed a discontinuous axonemal pattern in DasR cells compared to control cells (Fig. 1J). In fact, through super-resolution microscopy, we find that these Arl13B-positive fragments (Fig. 1K) are completely surrounded by Arl13B-containing membrane (Fig. 1L).

Next, we wanted to understand the molecular nature of the longer cilia phenotype. The kinesin Kif7 has been shown to control cilia length by organizing the cilia tip in coordination with the IFT-B particle IFT81 (He et al., 2014). In fact, the changes in cilia length observed in DasR cells are reminiscent of those seen in Kif7-deficient cells, suggesting that Kif7 could be a mediator of this phenotype. We found that in control cells, Kif7 localized to the ciliary base, along the cilium, and at the cilium tip, as previously described (He et al., 2014) (Fig. 2A). In contrast, DasR cells had a significant decrease in Kif7 localization to the axoneme and no Kif7 localization to the cilia tip (Fig. 2A) while total Kif7 levels remained unchanged (Fig. 2D). We also found that in control A204 cells, IFT81 localized along the axoneme and at the cilia tip. However, it was absent from the axoneme in DasR cells (Fig 2B). The microtubule plus-end-binding protein EB1, which localizes to centrioles and cilia tips (Pedersen et al., 2003), is also thought to play a role in cilia biogenesis (Schroder et al., 2011). We found that localization of EB1 was restricted to centrioles in control A204 cells, while in DasR cells, EB1 localized along the ciliary axoneme as well as the cilia tip (Fig. 2C, C’). Thus, control of cilia length and cilia tip compartment organization, as well as cilia transport appear compromised in DasR cells.

Kif7 inactivation has been shown to destabilize cilia (He et al., 2014). Thus, we tested the stability of cilia in DasR cells by subjecting them to cold treatment or the microtubule destabilizing agent nocodazole. Both treatments resulted in significantly shorter cilia in DasR cells compared to untreated controls (Supplementary Fig. 2D-E,F-G). Thus, elongated cilia in DasR cells are unstable. This is consistent with our observation that cilia in drug-resistant cells are less polyglutamylated (Supplementary Fig. 2C), a modification shown to regulate microtubule stability (O’Hagan et al., 2011).

Given these results, we hypothesized that Kif7 downregulation would promote cilia elongation and increase resistance to kinase inhibitors. Notably, downregulation of Kif7 in drug-sensitive control cells, although growth inhibitory in the absence of drug, significantly increased resistance to Dasatinib (Fig. 2E), and caused the expected increase in cilia length (Fig. 2F, G, H). Conversely, overexpression of Kif7 promoted a decrease in cilia length (Fig. 2I, J).

Because drug resistance often involves aberrant activation of compensatory pathways (many of which reside in or are controlled by cilia), we hypothesized that the observed changes in cilia would lead to misregulated cilia-dependent signaling. Activation of the evolutionarily conserved Hedgehog (Hh) pathway requires a functional cilium, and it is coordinately regulated at the body of the cilium and the cilium tip (Goetz et al., 2009; Huangfu and Anderson, 2005). Hh signaling is critical during development and for tissue maintenance in the adult (Hooper and Scott, 2005), and is aberrantly activated in some cancers, where it serves as a therapeutic target (Pak and Segal, 2016). The Hh pathway is activated when Hh ligands (e.g. Sonic Hh (Shh)) bind the ciliary localized transmembrane receptor Patched (PTCH). This promotes PTCH removal from the cilium and relieves inhibition of the key signal transducer Smoothened (SMO). SMO moves into the cilium and activates GLI transcription factors leading to Hh-specific target gene transcription (Goetz et al., 2009). Given that changes in cilia tip organization, KIF7 defects, and changes in ciliogenesis have been shown to disrupt the Hedgehog pathway (He et al., 2014), we hypothesized that cilia changes in resistant cells might affect Hh function. We first examined SMO recruitment to the cilium following Shh stimulation (Fig. 3A). Fluorescence intensity quantification showed increased SMO recruitment to cilia in DasR cells compared to control cells (Fig. 3A,B). To assess the functional relevance of this increase, we examined transcriptional targets of Hh-activation by RT PCR at steady state, and at different time points after addition of human Shh. We found significantly higher induction of the Hh target genes GLI1 and PTCH1 in response to either Shh (Fig. 3C, D) or the SMO agonist (SAG) (Supplementary Fig. 3) in DasR cells compared to A204 control cells, thereby confirming that the longer cilia observed in DasR cells support enhanced Hh pathway activation (Figure 3A-D). Consistently, lung cancer H2228 cells with acquired resistance to the ALK inhibitor NVP-TAE684, also showed increased Hedgehog pathway activation (seen as both a significant increase in ciliary localization of smoothened following receptor engagement (Fig.3H, I), and an increase in the levels of GLI1) compared to parental controls (Fig.3J). Additionally, we observed an increase in GLI2 levels in erlotinib-resistant HCC4006 cells compared to parental controls (Fig.3K).

Our results indicate that acquired resistance to kinase inhibitors is associated with the upregulation of a number of ciliogenesis pathways, and suggest that targeting cilia might be an effective strategy to overcome resistance. To test this hypothesis, we asked whether inhibition of ciliogenesis via knockdown of the centriole distal appendage protein SCLT1 (Tanos et al., 2013) or the IFT-B particle IFT88 (Pazour et al., 2000) could affect KIR cell viability. Notably, while in our 4 models of acquired drug resistance disrupting ciliogenesis did not significantly alter the cell cycle (Supplementary table 1), when combined with the appropriate kinase inhibitor (i.e. erlotinib, dasatinib, or NVP-TAE684) it significantly reduced viability in drug-resistant cells (Fig. 4A, Supplementary. Fig. 5S, Fig. 4B). Furthermore, IFT88 knockdown in DasR cells significantly reduced anchorage-independent growth (Fig. 4C).

**Figure 4.**
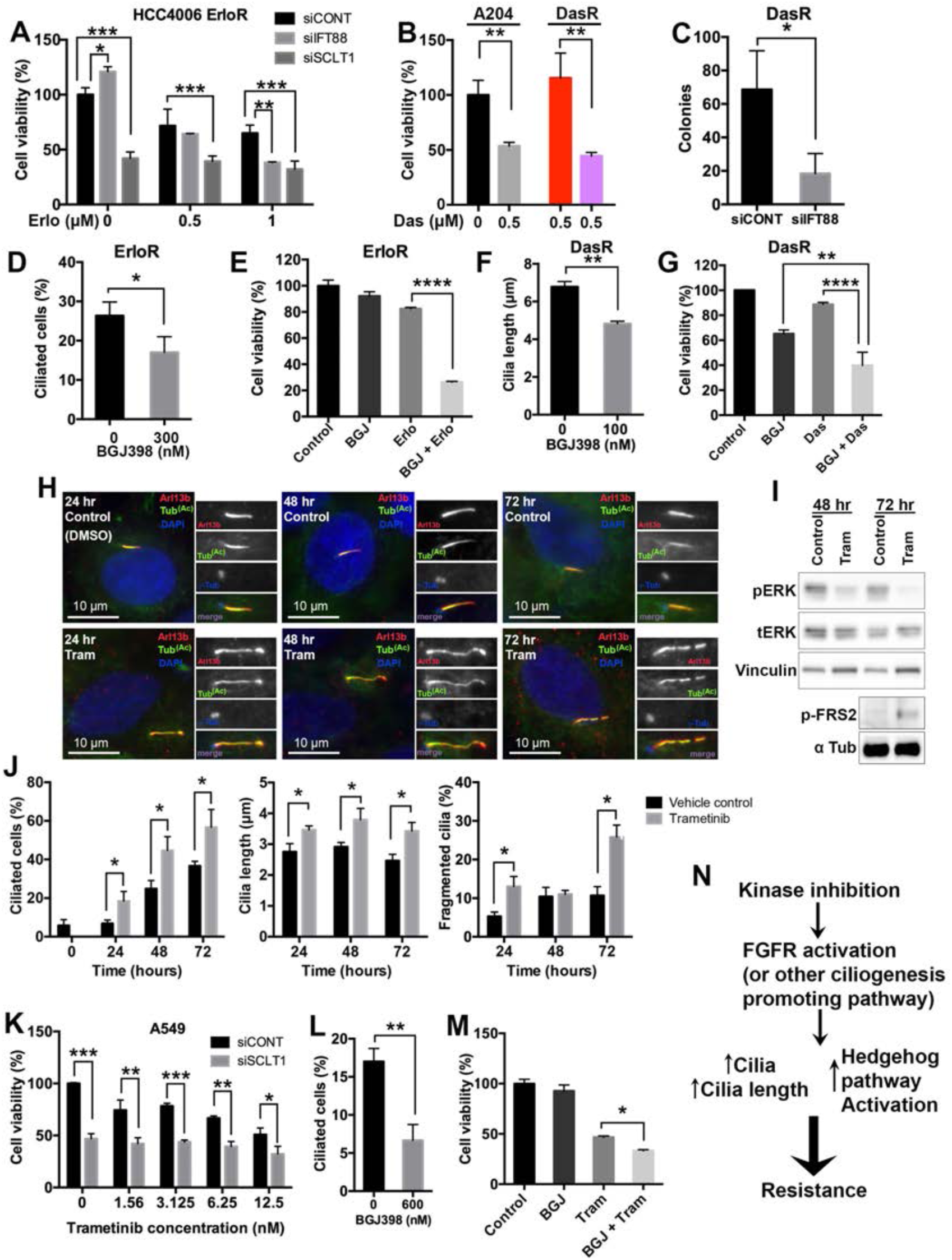
Ciliary pathways are key during the onset of kinase inhibitor resistance. (**A**) Cell viability (Cell titer Glo) in HCC4006 cells grown in erlotinib (indicated), were transfected with either control siRNA, IFT88 siRNA, or SCLT1 siRNA (indicated). Cell viability was normalized to siCONT DMSO control (0 μM) treated cells (n = 3); error bars represent s.d. p<0.05 (siControl compared to siIFT88 at 0 μM), p<0.0001 (siControl vs siSCLT1 at 0 μM), p<0.0008 (siControl vs siSCLT1 at 0.5 μM), p<0.006 (siControl compared to siIFT88 at 1 μM), p<0.0007 (siControl compared to siSCLT1 at 1 μM), Tukey’s multiple comparison test. (**B**) Cell viability in A204 cells, grown in the absence or the presence of dasatinib (indicated), and in DasR cells, after transfection with control siRNA or IFT88 siRNA (Robert et al., 2007). Cell viability is normalized to A204 DMSO control (0 μM) (n = 3). Error bars represent s.d. p<0.005 for A204 grown in 0 dasatinib compared to 0.5 μM dasatinib and p<0.006 for DasR siCONT compared to siIFT88, unpaired T test. (**C**) Soft agar colony formation in DasR cells transfected with control (non-targeting) siRNA or siRNA for IFT88 (Robert et al., 2007). Error bar bars represent s.d. p<0.03, unpaired T test, n = 3. (**D**) Cilia length of HCC40006 erlotinib resistant cells (ErloR) treated with or without the FGFR inhibitor BGJ398 for 72 hours. Note that after treatment with BGJ398 cilia length was reduced. n = 150, error bars represent s.d. p<0.04, unpaired T test. (**E**) Cell viability (Cell titer Glo) of ErloR grown in 1 μM erlotinib (Erlo) and 300 nM BGJ398. Note that combining both erlotinib and BGJ398 significantly reduced growth compared to erlotinib used as a single agent. Cell viability was normalized to DMSO control (n=3). Error bars represent s.d. p<0.0001, for an unpaired T test. (**F**) Cilia length of A204 dasatinib resistant cells (DasR) treated with or without BGJ398 for 24 hours in reduced serum conditions (5 % FBS). Note that after treatment with BGJ398 cilia length in DasR cells was reduced. n = 150, error bars represent s.d. p<0.0005, unpaired T test. (**G**) Cell viability (Cell titer Glo) of DasR cells treated with dasatinib (0.5 μM), the FGFR1 inhibitor BGJ398 (100 nM), or a combination of both. n = 3, cell viability is normalized to the DMSO control. Error bars represent s.d. p<0.004 (BGJ vs BGJ + Das), p<0.0001 (Das vs BGJ + Das), Tukey’s multiple comparison test. (**H**) A549 cells were treated with 50 nmol/L Trametinib (Tram) or DMSO (vehicle control) for the indicated times then fixed and stained with antibodies for acetylated tubulin (green), Arl13B (red), γ-tubulin (blue/inset) and DAPI (blue). Note that exposure to trametinib promoted a significant increase in cilia, cilia length and fragmentation. (**I**) Western blot showing A549 expression of phosphorylated ERK, total ERK, vinculin (loading control), phospho-FRS2 and α-tubulin (loading control) in the presence and absence of 50 nM trametinib (Tram). Note that after 48 and 72 hours of trametinib exposure pERK was reduced. Phospho-FRS2 expression increased after 72 hours. (**J**) Quantification of ciliated cells, cilia length and fragmentation shown in **H**. n = 150 cilia, error bars represent the s.d. p<0.05 unpaired T test. (**K**) Cell viability in A549 cells after treatment with the MEK inhibitor trametinib in control cells (siCONT) or upon down-regulation of the distal appendage protein SCLT1 (siSCLT1). Cell viability was normalized to siCONT DMSO control (0 nM) cells. n = 3, p<0.0001 for 0 nM, p<0.008 for 1.56 nM, p<0.0009 for 6,25 nM and p<0.003 for 12,5 nM, unpaired T test. (**L**) Cilia length of A549 cells treated with or without the FGFR inhibitor BGJ398 for 48 hours. Note that after treatment with BGJ398 cilia length of cells was reduced. n = 150, error bars represent s.d. p<0.003, unpaired T test. (**M**) Cell viability (Cell titer Glo) of A549 cells treated with trametinib (6.25 nM), BGJ398 (300 nM), or a combination of both. n = 3, cell viability is normalized to the DMSO control. Error bars represent s.d. p<0.02, for an unpaired T test. (**N**) Proposed model for up-regulation of ciliogenesis during kinase inhibitor resistance (KIR) acquisition.

Next, we interrogated the impact of pharmacological targeting of cilia function on drug resistance. First, we focused on the Hh pathway because it is upregulated in DasR cells, and because it has previously been implicated in drug resistance (Faiao-Flores et al., 2017). Interestingly, we found that treatment with Gant61, a small molecule inhibitor of the Hh pathway (Lauth et al., 2007) reduced viability in both DasR and control cells (Supplementary Fig.3E), highlighting the broad therapeutic potential of targeting cilia function in cancer. Second, we targeted fibroblast growth factor receptor (FGFR) because it has been previously shown to control ciliogenesis and cilia length (Neugebauer et al., 2009). Accordingly, we found that treatment of erlotinib-resistant HCC4006 cells with the FGFR inhibitor BGJ398 significantly reduced cilia formation (Fig. 4D), and more importantly, it re-sensitized these cells to erlotinib (Fig. 4E). We found similar results when we evaluated FGFR inhibition in A204 DasR cells (Fig. 4F-and NVP-TAE684-resistant H2228 cells (Supplementary Fig. 6), suggesting that inhibition of cilia regulators such as FGFR may represent a good therapeutic strategy to overcome drug resistance in a variety of contexts.

Finally, to assess the scope of our findings, we wanted to know whether cilia changes similar to those in our models of acquired resistance were associated with any instances of *de novo* drug resistance.

In NSCLC, KRAS is targeted by activating mutations in 20-30% of the cases. However, direct targeting of RAS has been challenging. One approach to target RAS function has been to inhibit components of the downstream MAPK pathway, including MEK. However, KRAS-mutant cells are largely refractory to these drugs. Two independent studies have found that in KRAS-mutant lung cancer cells, FGFR can mediate adaptive resistance to the MEK inhibitor, trametinib, (Kitai et al., 2016; Manchado et al., 2016).

We therefore hypothesized that MEK-inhibitor resistance in KRAS-mutant A549 cells would be accompanied by changes in ciliogenesis. Notably, we found that cilia number as well as cilia length were upregulated in A549 cells following trametinib treatment (Fig. 4H-J), although changes in Hedgehog pathway activation were difficult to assess due to the significantly high level of basal activity (Supplementary Fig 4J-K). Additionally, KRAS-mutant NCI-H23 and NCI-1792 lung cancer cells also showed upregulated ciliogenesis and increased Hedgehog pathway activation (Supplementary Fig 4, Table 1 in response to MEK inhibition. Furthermore, after 24 hours of drug treatment, the elongated cilia in A549 cells started to show evidence of fragmentation (Fig. 4H, J). Thus, release of terminal cilia-fragments might be a common feature of KIR cells independently of the molecular identity of the resistance pathway.

Importantly, inhibiting ciliogenesis in all 3 KRAS-mutant lines reduced their viability when combined with trametinib (Figure 4, Supplementary Fig. 5), while having no significant changes in cell cycle distribution (Supplementary table 1) on its own.

Furthermore, similar to our models of acquired kinase inhibitor resistance and consistent with previous reports (Kitai et al., 2016; Manchado et al., 2016), treatment with the FGFR inhibitor BGJ398 significantly reduced their viability in the presence of trametinib (Fig. 4L, M, Supplementary Figure 6).

These data support a model wherein inhibition of certain kinases leads to increased activation of FGFR (or other cilia promoting pathways), leading to enhanced ciliogenesis and concomitant hedgehog pathway activation, thus facilitating the generation of inhibitor insensitive survival signals (Fig. 4N). Cilia could thus function as a permissive platform for a number of drug resistance mechanisms with broad therapeutic implications.

## Discussion

Our work has uncovered ciliogenesis and cilia function as key biological processes that play permissive roles in the emergence of resistance to kinase inhibitors in cancer cells. Although a large number of tumors show a decrease in the number of cilia or fail to produce primary cilia, our data show that resistance to a variety of targeted therapies in several experimental models is characterized by an increase in the number and length of primary cilia and by cilia fragmentation. The significance of these fragments is beyond the scope of this study, but it is important to note that they will likely add to the complexity of tumor heterogeneity.

Consistent with the notion that aberrant cilia can alter oncogenic signaling, we find that the local abundance of a number of cilia-associated oncoproteins changes in dasatinib resistant cells. For example, DasR cells lose PDGFRα expression (Supplementary Fig. 1F, (Wong et al., 2016)), show a slight increase in FGFR1 (Supplementary Fig. 1G), and have increased ciliary localization of IGF-1R (Supplementary Fig. 1H).

Furthermore, cilia elongation in resistant cells coincides with a decrease of Kif7/IFT81 localization to cilia. In control cells, Kif7 is at the cilia tip, where it promotes microtubule plus-end catastrophe, thus creating a tip compartment for the enrichment of IFT81(He et al., 2014). In contrast, in DasR cells the Kif7-rich cilia tip compartment is lost, which explains the absence of IFT81. Thus, KIR cells have clearly defined molecular changes at the cilia tips. Additionally, we observed increased EB1 localization along the cilia and cilia-tips in KIR cells (Fig. 2C), which is suggestive of defective diffusion barrier control. Notably, downregulation of Kif7 in dasatinib-sensitive cells resulted in longer cilia and rendered these cells resistant to treatment.

Our results show that KIR cells have an enhanced response to Hedgehog pathway activation (Fig. 3, Supplementary Figure 3 and 4), and that DasR cells are sensitive to the Gli inhibitor Gant61 (Supplementary Fig. 3E), suggesting that drug resistance may be mediated by a critical effector of cilia-dependent signaling.

Interestingly, we and others have shown that resistance to both the MEK inhibitor trametinib and the tyrosine kinase inhibitor dasatinib can be mediated by activation of FGFR (Manchado et al., 2016) (Kitai et al., 2016) (Wong, Finetti et al. 2016) (this paper), a kinase known to regulate cilia length (Neugebauer et al., 2009). Notably, in all isogenic models studied, we found that treatment of KIR cells with an FGFR inhibitor not only restored kinase inhibitor sensitivity (Fig. 4, Supplementary fig 6), but also reduced cilia and/or cilia length (Fig. 4D, F, L). Similarly, targeting ciliogenesis through knockdown of the centriole distal appendage protein SCLT1 (Tanos et al., 2013) or the IFT particle IFT88 (Pazour et al., 2000) sensitized cells to the relevant kinase inhibitor.

This is in contrast to the lack of sensitizing activity of therapeutic agents that cause growth arrest in specific phases of the cell cycle (i.e. cisplatin, rapamycin, or doxorubicin), which suggests that the sensitizing effects of ciliogenesis inhibition in drug-resistant cells are unlikely to be attributed to cell cycle deregulation (Supplementary Fig. 7). It is also unlikely that drug-resistance-associated changes in cilia are caused by alterations in cell cycle-dependent signals (Plotnikova et al., 2009), given that drug-resistant cells did not show any significant differences in cell cycle distribution compared to their parental counterparts (Supplementary Table 1).

Aurora A kinase is thought to be a critical factor for cilia disassembly (Pugacheva et al., 2007). However, we found no evidence of changes in Aurora A kinase activation in any of our isogenic models (not shown).

Of note, we have also examined ciliation in an A549 isogenic model of acquired chemoresistance, and found that resistance to Cisplatin and Vinflunine is also associated with a significant increase in ciliogenesis (Table 1, Supplementary Figure 8), suggesting that cilia might be involved in resistance to a wide range of therapeutic agents.

In summary, our study shows for the first time that aberrant ciliogenesis could serve as a functional platform for a variety of cancer drug resistance mechanisms (both *de novo* and acquired), and provides rationale for a broad therapeutic strategy to overcome resistance in a variety of settings.

## Acknowledgements

We thank Carsten Janke (Institut Curie) for advice and reagents, Jacek Gaertik (University of Georgia), Kathryn Anderson (MSKCC) and Robert Blassberg (CRICK) for kindly sharing antibodies. Max Liebau (University of Cologne), Stephane Angers (University of Toronto) and Kathryn Anderson (MSKCC) for kindly donating Kif7 expression plasmids. Special thanks to Marc Fivaz and Fredrik Wallberg from the ICR Imaging core. We thank Igor Vivanco (ICR) and Tony Magee (Imperial College London) for critically reading this manuscript.

## Methods

### Cell culture

Cells were maintained in DMEM (A549, A204 and the dasatinib resistant sub-line DasR), and DME/F12 (HCC4006) containing 10 % FBS, 4 mM GlutaMax (Thermo Scientific, Waltham, MA, USA), 500 μg/ml Normocin (InvivoGen, San Diego, CA, USA), 100 units/ml penicillin and 100 mg/ml streptomycin (Thermo Scientific). 5 μM dasatinib (LC Labs, Woburn, MA, USA) was supplemented to DasR growth media. The erlotinib resistant HCC4006 sub-line was grown in the presence of 1 μM erlotinib (LC Labs). NCI-H23 and NCI-H1792 were maintained in RPMI containing 10 % FBS, 2 mM GlutaMax (Thermo Scientific), 500 μg/ml Normocin (InvivoGen), 100 units/ml penicillin and 100 mg/ml streptomycin (Thermo Scientific). NCI-H2228, PC9, A549 and the cisplatin, carboplatin and vinflunine resistant sub-lines were maintained in IMDM containing 10 % FBS, 2 mM GlutaMax (Thermo Scientific), 500 μg/ml Normocin (InvivoGen), 100 units/ml penicillin and 100 mg/ml streptomycin (Thermo Scientific). The NVP TAE resistant NCI-H2228 sub-line was supplemented with 0.5 μM NVP-TAE684 (Axon Medchem, Cat# Axon 1416). PC9 resistant sub-lines were cultured with 2 μM afatinib (Stratech Scientific, Cat# S1011-SEL) and 10 μM erlotinib (LC Labs). A549 resistant sub-lines were cultured in 2 μg/ml cisplatin (Cayman Chemical Company, Ann Arbor, MI, USA) and 10μg/ml carboplatin (Cayman Chemical Company). HEK-293T cells were maintained with DMEM containing 10 % FBS, 2 mM GlutaMax (Thermo Scientific), 100 units/ml penicillin and 100 mg/ml streptomycin (Thermo Scientific).

### Ciliogenesis experiments

To induce cilia formation cells were plated on to polylysine coated coverslips in 3.5-cm plates at 0.4 x 10^6^ cells per well, allowed to attach for 24 hours then serum starved for 48 hours. For A549, NCI-H23 and NCI-H1792, ciliogenesis experiments were carried out in the presence of serum, since trametinib proved to be toxic otherwise. To activate the hedgehog pathway, cells were serum starved for 24 hours prior to the addition of 5 μg/ml Shh-N (Peprotech, London, UK) or 100 nM SAG (Millipore, Darmstadt, Germany). For cilia stability experiments, cells were either incubated in 4 °C culture media or treated with 10 μM nocodazole (Sigma-Aldrich, St. Louis, Missouri, USA).

### Immunofluorescence

Cells were fixed in 4 % paraformaldehyde for 10 min at room temperature, for the following antibodies: mouse anti-α-tubulin (1:200, YL1/2: Bio-Rad MCA77G); mouse anti-acetylated tubulin (1:2000, 6-11B-1; Sigma T7451); rabbit anti-Arl13B (1:500, Proteintech 17711-1-AP); mouse anti-EB1 (1:250, BD biosciences 5/EB1); mouse anti-centrin (1:500, 3E6; Abnova H00001070-M01); rabbit anti-detyrosinated α-tubulin (1:100, Abcam ab48389), rabbit anti-IGF-1Rβ (1:250, C-20; Santa Cruz sc-713), mouse anti-polyglutamylated tubulin (1:200, GT335; Adipogen AG-20B-0020) and rabbit anti-SMO (a kind gift by Kathryn Anderson, 1:500). An additional fixation step of 20 min in cold methanol was used for rabbit anti-IFT88 (1:500, Proteintech 13967-1-AP) and mouse anti-γ-tubulin (1:500; TU-30; Santa Cruz sc-51715). For antibodies against Kif7 (1:500, rabbit polyclonal, kind gift from Kathryn Anderson’s lab) and rabbit anti-IFT81 (1:200, Proteintech 11744-1-AP), cells were first permeabilized for 2 min in PTEM buffer (20 mM PIPES (pH 6.8), 0.2% Triton X-100, 10 mM EGTA, and 1 mM MgCl2) followed by fixation in cold methanol for 20 mins. After fixation, cells were permeabilized for 5 min in 0.1% Triton X-100 in PBS, then blocked with 3% (w/v) bovine serum albumin in PBS, 0.1% Triton X-100 for 5 min. Primary antibodies were diluted in blocking solution and incubated for 1h followed by 3 washes with PBS, 0.1% Triton X-100. After that, goat secondary antibodies conjugated to either Alexa Fluor 488, 594 or 680 (1:500 dilution; Thermo Scientific) were incubated for 1h followed by 3 washes and incubation with DAPI (Thermo Scientific).

### Image acquisition and analysis

Fluorescent images were acquired on an upright microscope (Axio Imager M2, Zeiss) equipped with 100x oil objectives, 1.4 NA, a camera (ORCA R2, Hamamatsu Photonics) and a computer with image processing software (Zen). Images were quantified for pixel density and cilia length using ImageJ and Matlab and assembled into figures using Photoshop (CS5, Adobe). 3D structured illumination images were acquired using an SR1 Elyra PS1 microscope (Zeiss), images were processed using the ImageJ plugin SIMcheck to remove artifacts and 3D videos were made using the Volocity software (PerkinElmer). For Matlab quantifications, we used a custom-written script to quantify fluorescent intensity profiles along cilia. Cilia were segmented in a user-interactive manner using the improfile function from Matlab. Improfile retrieves the intensity values of pixels along a multiline path defined by the user. Acetylated tubulin was used to define the cilium path and intensity profiles were retrieved from channels of interest for Kif7, IFT81 and EB1. To reduce noise, we measured the average fluorescent intensity of 3 pixels (above, on and below the path) for each position along the cilium. To compare intensity profiles along cilia of different lengths we divided cilium length into 10 bins and extracted the average fluorescent intensity for each bin. The script is available upon request.

### Western Blots

Cells were lysed in RIPA buffer (Sigma-Aldrich) supplemented with protease and phosphatase inhibitors (Thermo Scientific) on ice. Lysates were sonicated and cleared by centrifugation at 12000g at 4 °C for 30 mins. Samples were separated by SDS PAGE on 3 - 8 % polyacrylamide gradient gels followed by transfer to nitrocellulose membranes. Membranes were probed with primary antibodies against mouse anti-Erk (1:1000; 3A7; Cell Signaling 9107), rabbit anti-phospho-Erk (1:1000; Cell Signaling 9101), rabbit anti-Met (1:1000; D1C2; Cell signaling 8198), rabbit anti-phospho-Met (1:1000; D26; Cell Signaling 3077), rabbit anti-EGFR (1:1000; D38B1; Cell Signaling 4267), rabbit anti-phospho-EGFR (1:1000; D7A5; Cell Signaling 3777), rabbit anti-Gab1 (1:1000; Cell Signaling 3232), rabbit anti-phospho-Gab1 (1:1000; C32H2; Cell Signaling 3233), mouse anti-GLI1 (1:750; L42B10; Cell Signaling 2643), rabbit anti-phospho-FRS2-α (1;500; Cell Signaling 3864), rabbit anti-GLI2 (1:500; H-300; Santa Cruz sc-28674), mouse anti-FLAG (1:1000; M2; Sigma F1804), rabbit anti-IGF-1Rβ (1:250; C-20; Santa Cruz sc-713), rabbit anti-Kif7 (1:500), rabbit anti-SCLT1 (1:500; Sigma HPA036560) mouse anti-β-Actin (1:2000; AC-74; Sigma A5316), mouse anti-Aurora-A Kinase (1:500; 4/IAK1; BD Bioscience 610939), rabbit anti-PDGFRα (1:500; D1E1E; Cell Signaling 3174), rabbit anti-FGFR1 (1:1000; Abcam EPR806Y), rabbit anti-IFT81 (1:500; Proteintech 11744-1-AP) mouse anti-EB1 (1:500; BD biosciences 5/EB1), rabbit anti-IFT88 (1:500; Proteintech 13967-1-AP), mouse anti-α-tubulin (1:1000; 236-10501; A11126 Thermo Scientific); mouse anti-vinculin (1:2000; hVIN-1; Sigma V9131) and mouse anti-transferrin receptor (1:500; H68.4; Thermo Scientific 136890), secondary antibodies were HRP conjugated rabbit or mouse anti-IgG antibodies (1:2000; Cell Signaling).

### siRNA gene knockdown

siRNA mediated IFT88 knockdown was carried out using two pooled sequences 5’-CGACUAAGUGCCAGACUCAUU-3’ and 5’-CCGAAGCACUUAACACUUA-3’ previously described (Robert et al., 2007), when indicated, or a SMARTpool ON-TARGETplus siRNA (GE Dharmacon, Lafayette, CO, USA). SCLT1 knockdown was achieved using a siGENOME Smartpool siRNA (GE Dharmacon). Kif7 was downregulated using a SMARTpool ON-TARGETplus siRNA (GE Dharmacon). Cells were transfected with Lullaby (Oz Biosciences, San Diego, CA, USA) (three sequential transfections) or Lipofectamine RNAimax for the Smartpool (two sequential transfections). Non-targeting (control) siRNA was purchased from Qiagen (#1027281).

### Plasmids and transfections

The human full-length FLAG-Kif7 construct was a kind gift from Dr Max Liebau, University of Cologne, Germany. Cells were transfected with Lipofectamine 3000 (Thermo Scientific).

All lentiviruses were generated by transient co-transfection of 293T cells with packaging and envelope vectors using PEI transfection reagent. The TRIPZ inducible human shIFT88 plasmid (GE Dharmacon), was used for stable IFT88 gene knockdown. H23 cells were selected for using 2 μg/ml puromycin

### Cell viability assays

4000 cells/well (2000 cells/well for A204/DasR) were seeded in to a 96 well plate (Greiner Bio-One, Kremsmunster, Austria) and incubated for 24 hours at 37 Celsius degrees, 5% CO_2_. After that, media (5 % FBS) containing drugs or vehicle controls was added to the cells and incubated for an additional 72 hours. Cell viability was measured using Cell Titer Glo (Promega), using a Victor X5 2030 Multilabel plate reader (Perkin Elmer). Cisplatin was obtained from Cayman Chemical Company, doxorubicin from LC Labs and rapamycin from Calbiochem (San Diego, CA, USA).

### Cell cycle analysis

To determine the cell cycle distribution, DNA content was assessed using propidium iodide (PI) staining. Cells were trypsinised and fixed in ice cold 70 % ethanol then stained with 20 μg/ml PI and 100 μg/ml RNAase A for 30 mins. Samples were run using a BD LSR II flow cytometer (BD Biosciences) and FlowJo to analyse results.

### Hedgehog pathway quantitative Reverse Transcription PCR

RNA was extracted using RNA mini kit (Thermo Scientific). Primers and TaqMan probes for detection of human Tata binding protein (TBP), GLI1, and PTCH1 were purchased as Assays-on-Demand from Applied Biosystems (TBP: Hs00427620_m1, GLI1: Hs01110766_m1, PTCH1: Hs00181117_m1). SuperScript III Platinum One-Step qRT-PCR System (Invitrogen) was used for the qPCR (PCR protocol: 15 min 50°C, 2 min 95°C, 30-50x 15sec 95°C and 1 min 60°C). The amount of amplicon generated during the PCR was measured using a QuantStudio 6 Flex Real-Time PCR System (Applied Biosystems). Each sample was run in triplicate; controls without reverse transcriptase gave no signal in all samples.

### Soft agar assay

Each well of a 6 well dish was coated with 1-ml base layer containing 0.6% agar (Sigma-Aldrich). Cells were dissociated and filtered through 30 *μ*m filter and sub-cultured by layering 1 x 10^4 viable cells in 1.5 ml culture medium (5 % FBS) containing 0.3% agar over replicate base layers. An upper layer of 2ml culture medium (5 % FBS) was applied to each well and changed every 3 days. Colonies were counted using Gelcount (Oxford Optronix).

### Statistical tests

Statistical analyses and samples sizes are specified in the figure legends. The error bars indicate either standard deviation or standard error.

---

**Supplementary figure 1.**
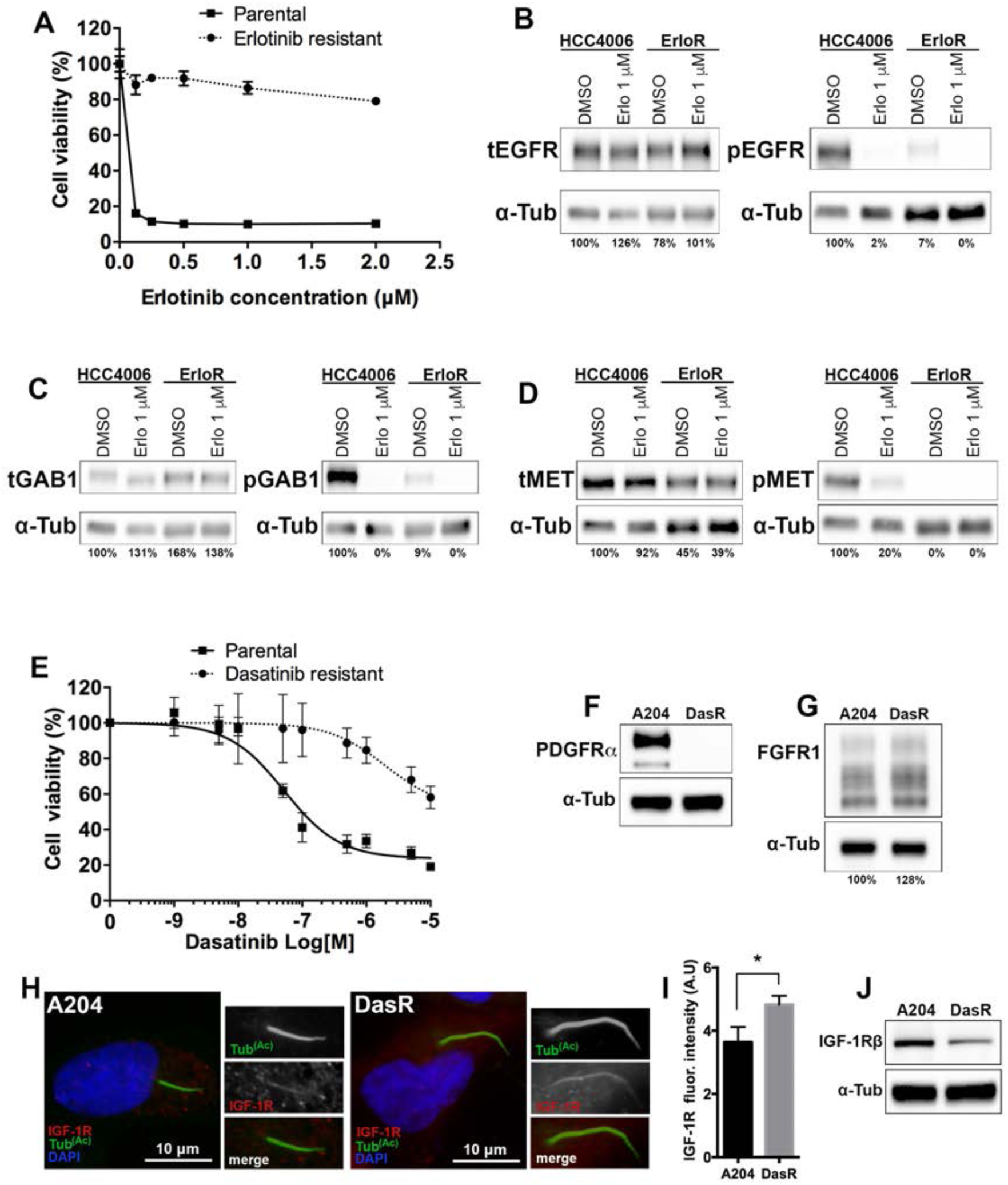
Molecular characterization of parental and kinase inhibitor resistant counterparts. (**A**) Dose response for erlotinib resistant cells. HCC4006 and the erlotinib resistant subline were treated with a range of concentrations of erlotinib to determine growth response curves. Cell viability was normalized to DMSO control (n=3). (**B, C, D**) Western blots showing HCC4006 parental and erlotinib resistant subline (ErloR) expression of total and phosphorylated: EGFR (**B**), GAB1 (**C**) and MET (**D**). α-tubulin (indicated) was used as a loading control. Cells were treated with or without erlotinib (1 μM) for 6 hours. (**E**) Dose response for dasatinib resistant cells. A204 and the dasatinib resistant subline (DasR) were treated with a range of dasatinib concentrations to determine growth response. Cell viability was normalized to DMSO control (n=3). (**F, G**) Western blots showing PDGFRα (**F**) and FGFR1 (**G**) of A204 and DasR cells (indicated). Note that DasR cells have no PDGFRα expression and a slight increase in FGFR1 expression. (**H**) DasR cells show increased ciliary localization of IGF-1Rβ compared to control cells. A204 (left) or DasR cells (right) were serum starved for 48 hours to induce ciliogenesis. After fixation, cells were stained with acetylated tubulin (green), IGF-1Rβ (red) and DAPI (DNA). (**I**) Quantification of IGF-1Rβ cilia fluorescence intensities shown in **H**. Fluorescence intensities were normalized to background camera fluorescence intensity. n = 150 cilia, error bars represent s.d. p<0.03, unpaired T test. (**J**) Western blot showing total expression levels of IGF-1Rβ (upper panel, indicated) and loading controls (lower panel) in A204 and DasR cells.

**Supplementary figure 2.**
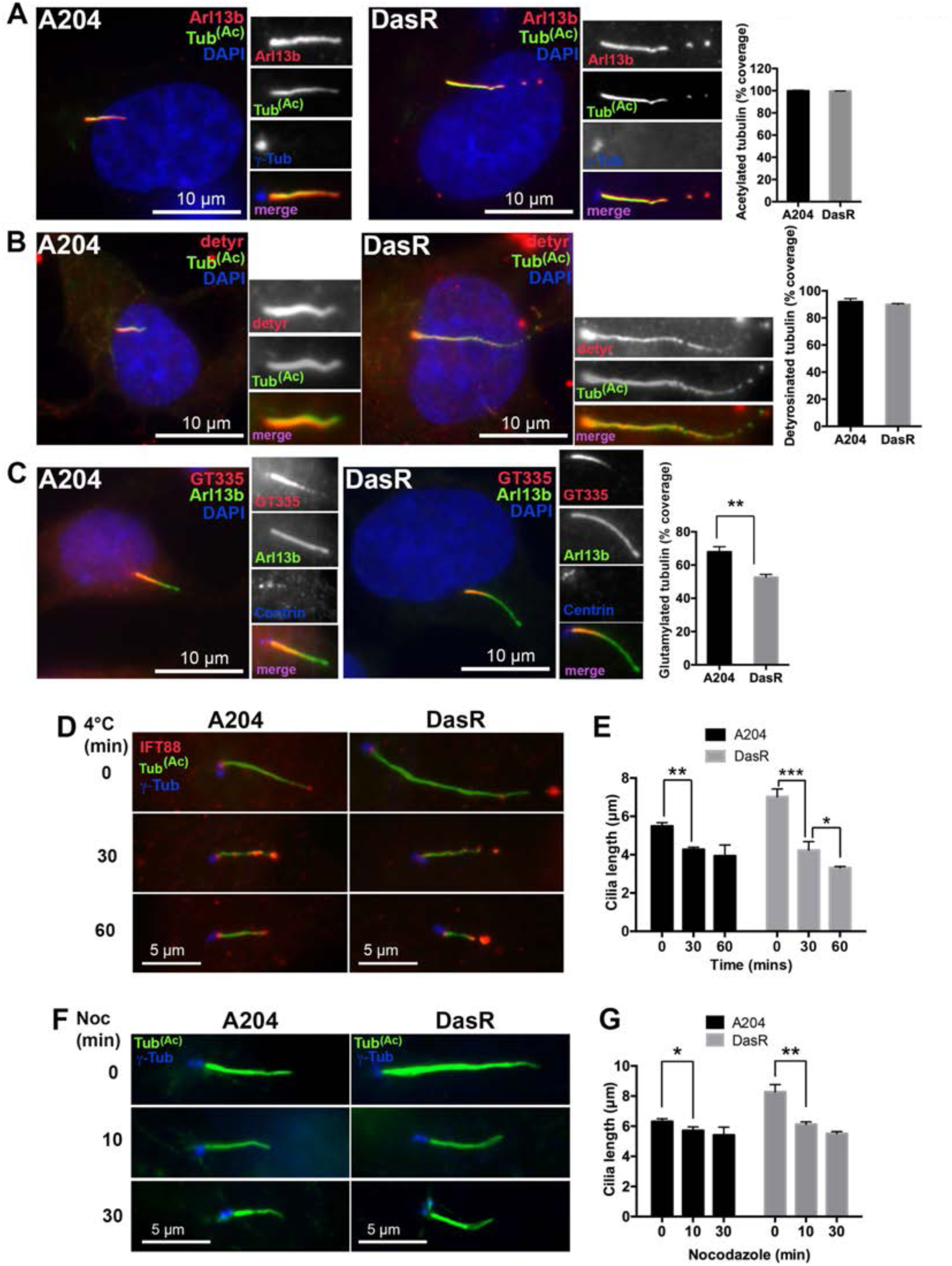
Kinase inhibitor resistant cells show decreased tubulin glutamylation along the axoneme and increased cilia instability. (**A, B, C**) Control (left panels) or dasatinib resistant cells (right panels) were serum starved for 48 hours to induce cilia formation, then fixed and stained with antibodies for Arl13b to mark the ciliary membrane (red), together with antibodies for different post-translational modifications (shown in green) including acetylated tubulin (**A**) detyrosinated tubulin (**B**) and glutamylated tubulin (GT335) (**C**). Centrioles are marked in blue (insets) with γ-tubulin or centrin (indicated) and DAPI is shown in blue. Graphs on the right show a quantitative analysis of the results for **A**, **B** and **C**, expressed as a function of total cilia length. n = 150, error bars represent s.d. p<0.01 unpaired T test. Note that DasR cilia show less polyglutamylated tubulin (GT335) along the axoneme compared to parental A204 cells. (**D**) Time course of cilia retraction in response to cold treatment (4°C) in A204 and DasR cells (indicated). Acetylated tubulin is shown in green, IFT88 in red and γ-tubulin in blue. (**E**) Quantification of cilia length in response to cold treatment for the experiment shown in **D**. Cilium length was measured using acetylated tubulin, n = 150. Error bars represent the s.d. p<0.003 between A204 at 0 and 30 mins, p<0.0001 between DasR at 0 and 30 mins and p <0.02 between DasR at 30 and 60 mins, Tukey’s multiple comparison test. (**F**) Time course of cilia retraction in response to nocodazole (10 μM) in A204 (left) and DasR cells (right). Acetylated tubulin staining marks primary cilia (green) and γ-tubulin marks centrioles (blue). (**G**) Cilium length was measured using acetylated tubulin staining from the time course shown in **F**. n = 150 cilia, error bars represent the s.d. p<0.02 for A204 0 and 30 mins and p<0.0001 DasR 0 and 30 mins, Tukey’s multiple comparison test. Note that DasR cells have an increased rate of cilia shortening in response to nocodazole.

**Supplementary figure 3.**
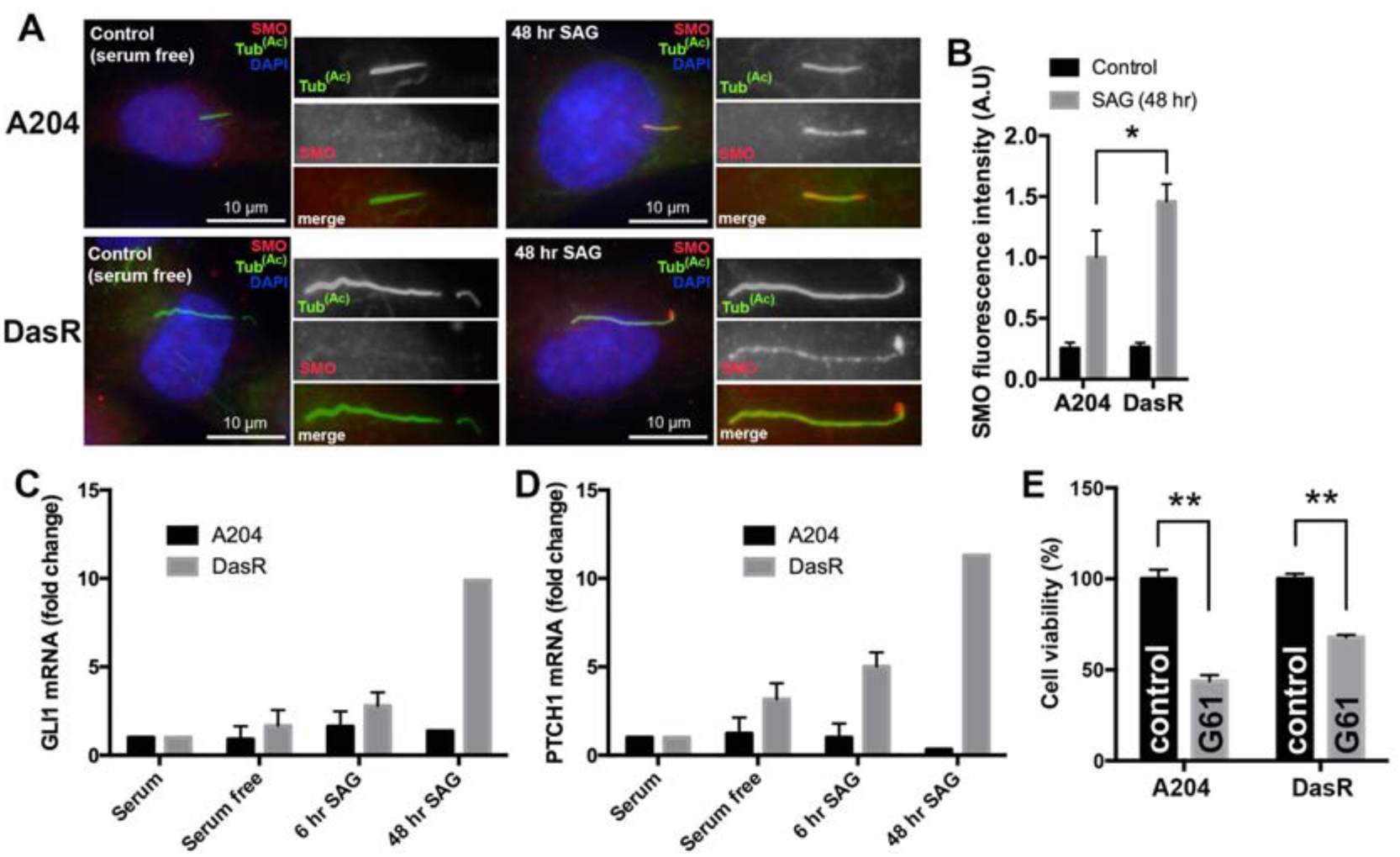
Upon SAG activation dasatinib resistant cells recruit more SMO into the cilium and activate the Hh pathway stronger and more prolonged compared to non-resistant control cells. (**A**) A204 cells (top panel), or a dasatinib resistant (DasR) subline (lower panel) were serum starved for 24 hours and either left in serum-free media for an additional 48 hours (left) or treated with SAG (100 nM) for the same amount of time (right). Cells were then fixed and stained with antibodies for acetylated tubulin to mark cilia (green), SMO (red) and with DAPI (blue) to mark DNA. (**B**) Quantification of SMO cilia fluorescence intensities for the experiment shown in **A**. Note the increased SMO fluorescence intensity in DasR compared to A204. Fluorescence intensity was normalized to surrounding fluorescence, n = 150, error bars represent s.d. p<0.04, unpaired T test. (**C, D**) Quantitative polymerase chain reaction (qPCR) showing fold change (relative to no serum starvation) mRNA levels of GLI1 (**C**) and PTCH1 (**D**) in A204 and DasR cells. Note the fold change of both *GLI1* (**C**) and *PTCH1* (**D**) is increased in DasR cells compared to A204 at all time points. GLI1 and PTCH1 mRNA values are normalized to TATA box–binding protein (*TBP*) mRNA values, fold change calculated by comparing to mRNA levels prior to serum starvation; n = 3 (0-6h). (**E**) Cell viability of A204 and DasR cells (indicated), in normal media (black columns) or with the addition Hh pathway inhibitor Gant61 (G61) (2.5 μM) (grey columns). Cell viability was normalized to DMSO control treated cells. p<0.01 unpaired T test.

**Supplementary figure 4.**
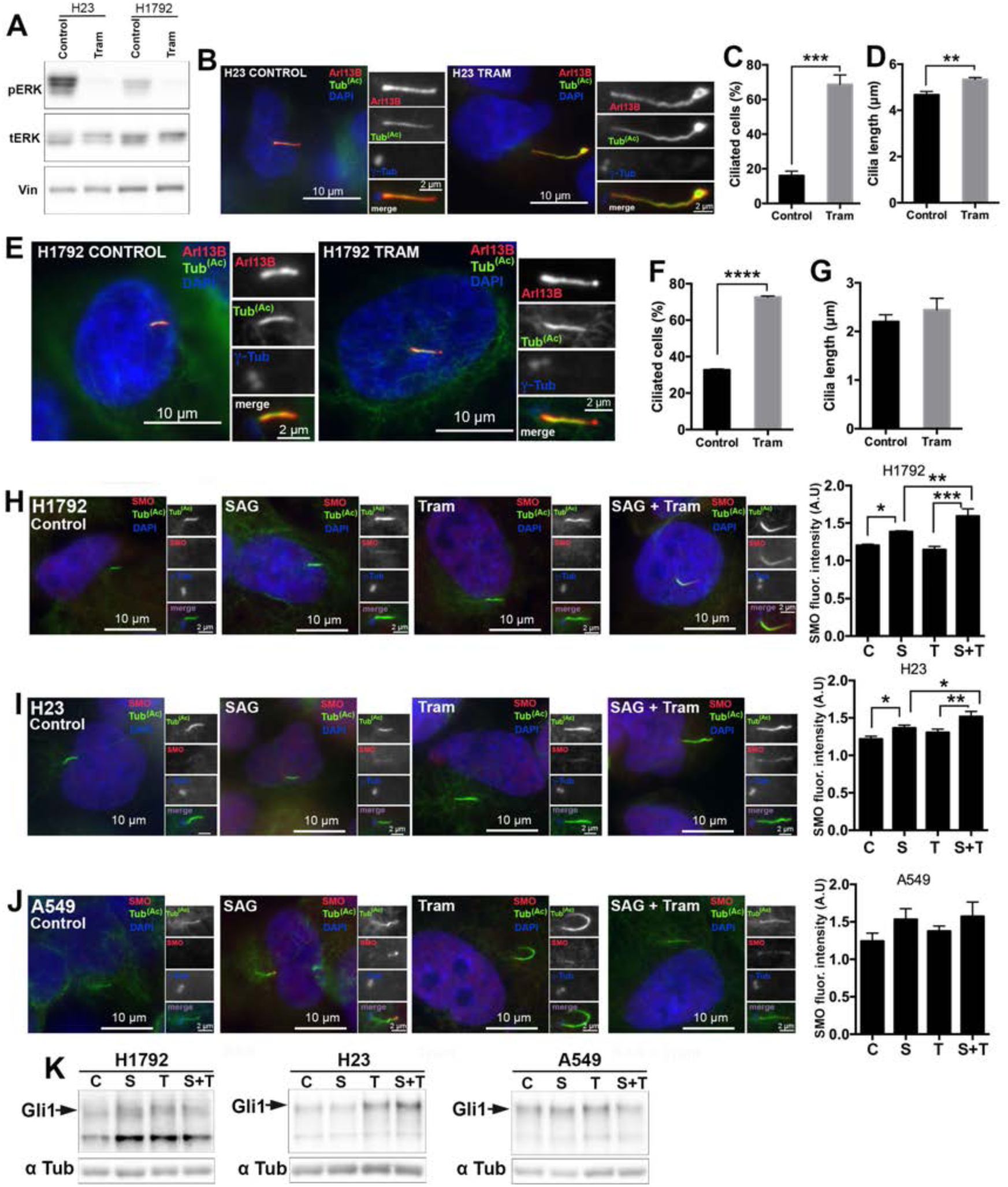
De novo drug resistance in patient-derived tumor cell lines shows increased cilia frequency, cilia length and hedgehog pathway activation. (**A**) Western blots showing NCI-H23 and NCI-H1792 expression of pERK in absence or presence of 50 nM trametinib (48 hrs). (**B**) NCI-H23 cells treated with 50 nM Trametinib or DMSO (control) for 48hrs then fixed and stained with antibodies for acetylated tubulin (green), Arl13B (red), γ-tubulin (blue/inset) and DAPI (blue). Note that exposure to trametinib promoted an increase in cilia length. (**C, D**) Quantification of ciliated cells (**C**) and cilia length (**D**) in **B**. n = 300 cells (**C**), n = 150 cilia (**D**), error bars represent the s.d. p<0.0001 (**C**) and p<0.003 (**D**), for an unpaired T test. (**E**) NCI-H1792 cells treated with 50 nM Trametinib or DMSO (control) for 48hrs then fixed and stained with antibodies for acetylated tubulin (green), Arl13B (red), γ-tubulin (blue/inset) and DAPI (blue). (**F, G**) Quantification of ciliated cells (**F**) and cilia length (**G**) shown in **E**. n = 300 cells (**F**), n = 150 cilia (**G**), error bars represent the s.d. p<0.0001 (**F**), for an unpaired T test. (**H**) NCI-H1792 cells were treated either with DMSO control (C), 100 nM SAG (S), 50 nM trametinib (T) or a combination of SAG and trametinib (S+T) for 48 hours. Cells were then fixed and stained with antibodies for acetylated tubulin to mark cilia (green), SMO (red) and with DAPI (blue) to mark DNA. Quantification of SMO cilia fluorescence intensities is shown on the right. Note the combination of trametinib and SAG increases SMO cilia fluorescence compared to SAG alone. Fluorescence intensity was normalized to surrounding fluorescence, n = 150, error bars represent s.d. p<0.02 (C vs S), p<0.0001 (T vs S+T), p<0.007 (S vs S+T), Tukey’s multiple comparison test. (**I**) NCI-H23 cells were treated either with DMSO control (C), 100 nM SAG (S), 50 nM trametinib (T) or a combination of SAG and trametinib (S+T) for 48 hours. Cells were then fixed and stained with antibodies for acetylated tubulin to mark cilia (green), SMO (red) and with DAPI (blue) to mark DNA. Quantification of SMO cilia fluorescence intensities is shown on the right. Note the combination of trametinib and SAG increases SMO cilia fluorescence compared to SAG alone. Fluorescence intensity was normalized to surrounding fluorescence, n = 150, error bars represent s.d. p<0.03 (C vs S), p<0.004 (T vs S+T), p<0.03 (S vs S+T), Tukey’s multiple comparison test. (**J**) A549 cells were treated either with DMSO control (C), 100 nM SAG (S), 50 nM trametinib (T) or a combination of SAG and trametinib (S+T) for 48 hours. Cells were then fixed and stained with antibodies for acetylated tubulin to mark cilia (green), SMO (red) and with DAPI (blue) to mark DNA. Quantification of SMO cilia fluorescence intensities is shown on the right. Fluorescence intensity was normalized to surrounding fluorescence, n = 150, error bars represent s.d. (**K**) Western blots showing GLI1 expression for H1792, H23 and A549 cells after 48hrs of DMSO (control) (C), SAG (S), Trametinib (T) or SAG and Trametinib (S+T) exposure.

**Supplementary figure 5.**
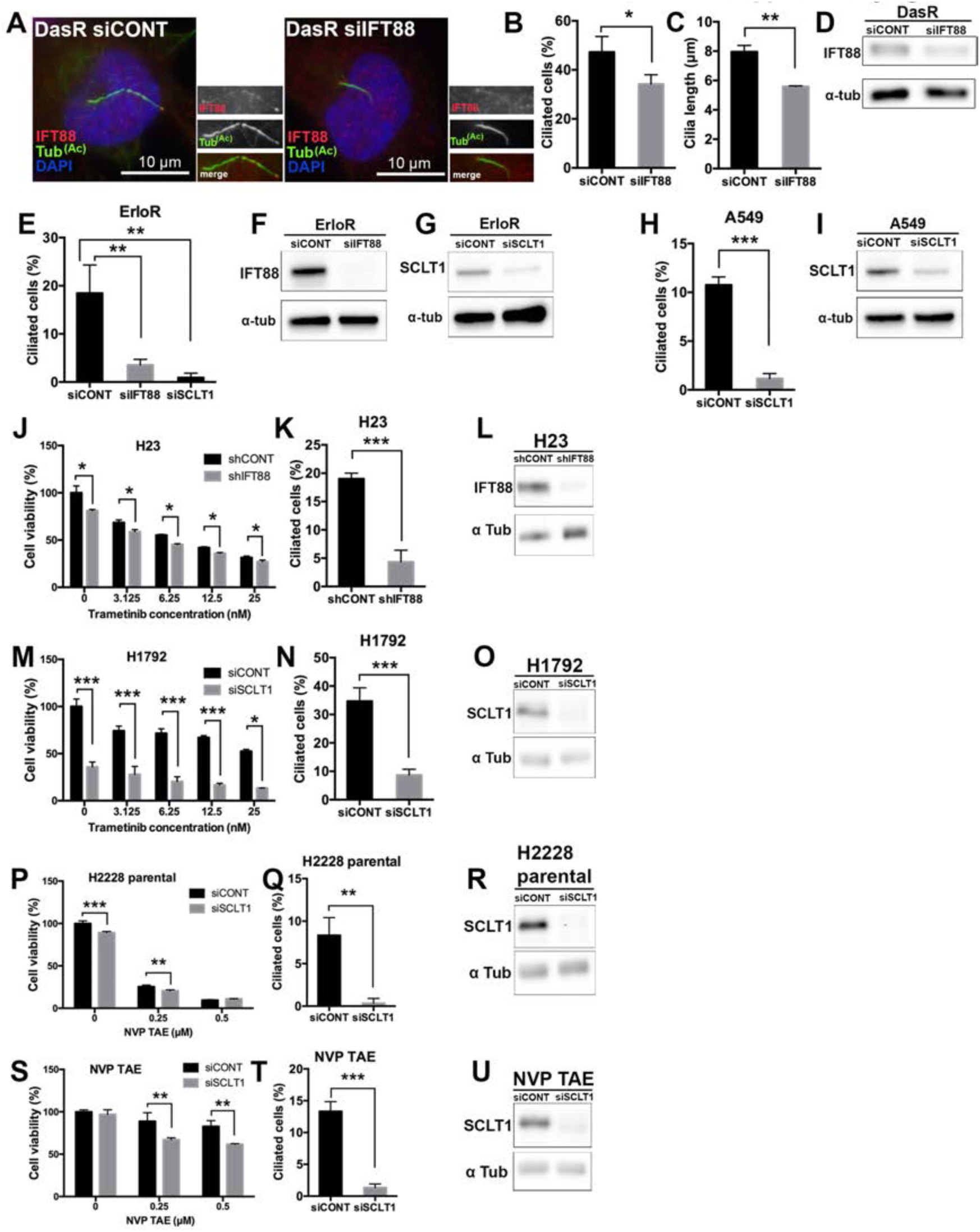
IFT88 and SCLT1 downregulation disrupt ciliogenesis in kinase inhibitor resistant cells. (**A**) DasR cells transfected with an IFT88 siRNA (siIFT88) (Robert et al., 2007) had reduced ciliated cells and cilia length compared to cells treated with a control siRNA (siCONT). DasR cells were serum starved for 48 hours to induce ciliogenesis, then fixed and stained with antibodies for acetylated tubulin (green), IFT88 (red) to mark cilia, and with DAPI (blue). (**B, C**) Quantification of percent ciliated cells (n=300), and cilia length (n=150) for the experiment shown in **A**. Error bars represent s.d. p<0.04 (for **B**), p<0.0008 (for **C**) unpaired T test. (**D**) Western blot showing IFT88 expression in DasR cells transfected with control siRNA or IFT88 siRNA (indicated) for the experiment in **A**, **B**, **C**. (**E**) Cilia quantification in HCC4006-Erlotinib resistant subline (ErloR) transfected with either control siRNA, siRNA for IFT88 (siIFT88) (Smartpool) or SCLT1 siRNA (siSCLT1) (Smartpool). Note that in both cases cilia frequency is significantly decreased compared to control siRNA. n = 300, error bars represent s.d. p<0.003 (siCONT vs siIFT88) and p<0.005 (siCONT vs siSCLT1). (**F**, **G**) Western blots showing IFT88 (**F**) or SCLT1 (**G**) expression of ErloR cells, transfected with control siRNA and either IFT88 siRNA (**F**) or SCLT1 siRNA (**G**) for the quantification shown in **E**. (**H**) A549 cells transfected with siRNA for SCLT1 (siSCLT1) had reduced ciliated cells compared to an siRNA control (siCONT). n = 300, error bars represent s.d. p<0.0001. (**I**) Western blot showing SCLT1 expression in A549 cells transfected with control siRNA or SCLT1 siRNA (indicated) for the experiment shown in **H**. (**J**) Cell viability in H23 cells after treatment with the MEK inhibitor trametinib in control cells (shCONT) or upon down-regulation of IFT88 with an inducible IFT88 shRNA (shIFT88). Cell viability was normalized to shCONT (DMSO). n = 4, p<0.003 for 0 nM, p<0.0001 for 1.56 nM, p<0.002 for 3.125 nM, p<0.0001 for 6,25 nM, p<0.0001 for 12.5 nM and p<0.001 for 25 nM, unpaired T test. (**K**) Quantification of percent ciliated cells (n=300) for the experiment shown in **J**. Error bars represent s.d. p<0.0005, unpaired T test. (**L**) Western blot showing IFT88 expression in H23 cells infected with shCONT or shIFT88 (indicated) for the experiment shown in **J**. (**M**) Cell viability in H1792 cells after treatment with the MEK inhibitor trametinib in control cells (siCONT) or upon down-regulation of SCLT1 (siSCLT1). Cell viability was normalized to siCONT (DMSO), p<0.0004 for 0 nM, p<0.002 for 3.125 nM, p<0.0003 for 6,25 nM, p<0.0001 for 12.5 nM and p<0.0001 for 25 nM, unpaired T test. (**N**) Quantification of percent ciliated cells (n=300) for the experiment shown in **M**. Error bars represent s.d. p<0.002, unpaired T test. (**O**) Western blot showing SCLT1 expression in H1792 cells transfected with siCONT or siSCLT1 (indicated) for the experiment shown in **M**. (**P**) Cell viability in H2228 parental cells after treatment with NVP TAE in control cells (siCONT) or upon down-regulation of SCLT1 (siSCLT1). Cell viability was normalized to siCONT (DMSO), n = 3, p<0.0001 for 0 μM, and p<0.004 for 0.25 μM, unpaired T test. (**Q**) Quantification of percent ciliated cells (n=300) for the experiment shown in **P**. Error bars represent s.d. p<0.004, unpaired T test. (**R**) Western blot showing SCLT1 expression in H2228 cells transfected with siCONT or siSCLT1 (indicated) for the experiment shown in **P**. (**S**) Cell viability in H2228 NVP TAE resistant cells after treatment with NVP TAE in control cells (siCONT) or upon down-regulation of SCLT1 (siSCLT1). Cell viability was normalized to siCONT (DMSO). n = 3, p<0.0006 for 0.25 μM, and p<0.006 for 0.5 μM, unpaired T test. Note that in the absence of cilia, NVP-TAE resistant cells become more sensitive to the inhibitor. (**T**) Quantification of percent ciliated cells (n=300) for the experiment shown in **S**. Error bars represent s.d. p<0.0003, unpaired T test. (**U**) Western blot showing SCLT1 expression in H2228 cells transfected with siCONT or siSCLT1 (indicated) for the experiment shown in **S**.

**Supplementary figure 6.**
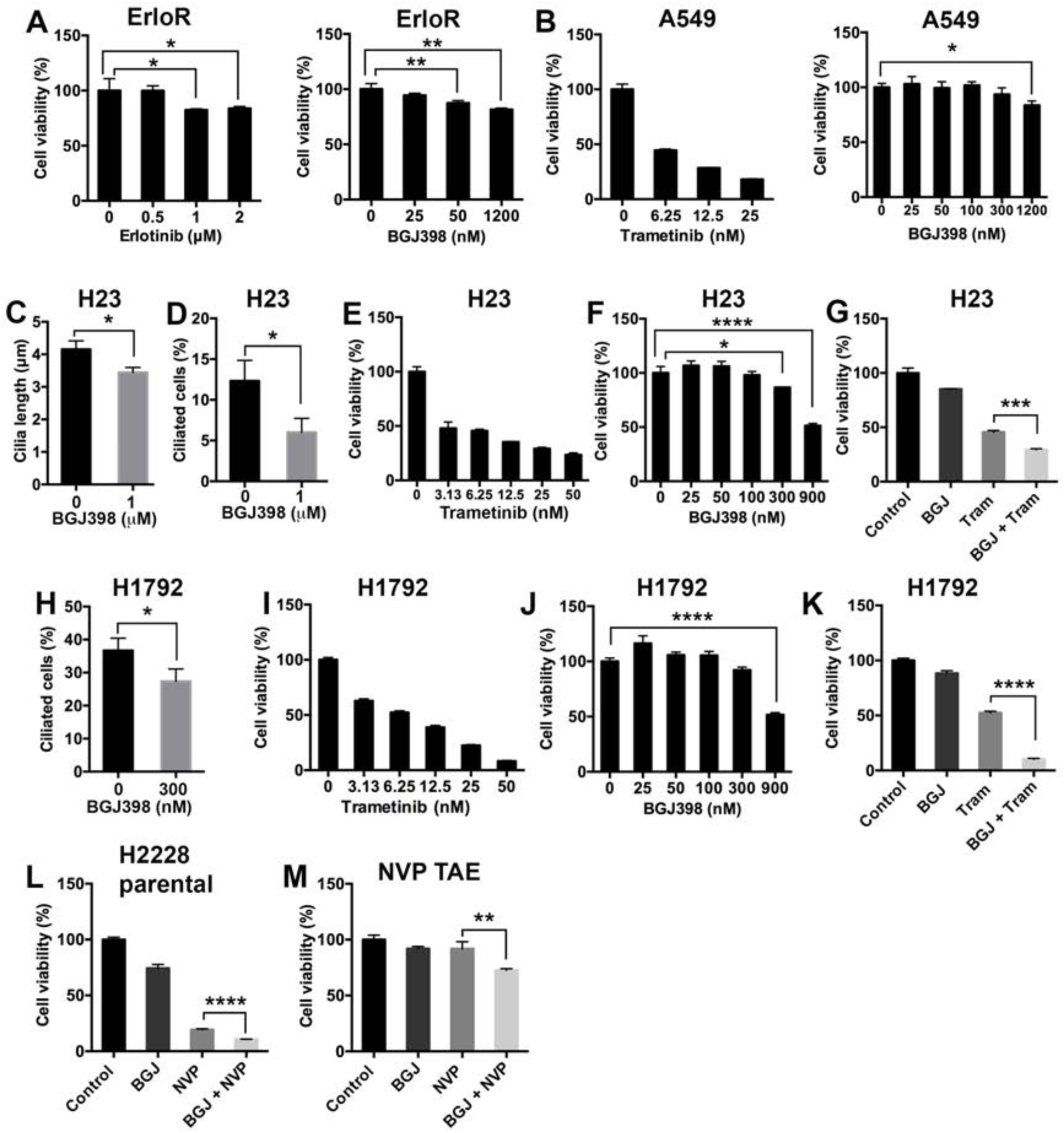
Targeting FGFR sensitises cells to kinase inhibitors. (**A**) Cell viability (Cell titer Glo) of the HCC4006 erlotinib resistant subline (ErloR) grown in a range of erlotinib (left graph) and the FGFR inhibitor BGJ398 (right graph) concentrations. Cell viability was normalized to DMSO control (n=3). Error bars represent s.d. p<0.03 (0 compared to 1 μM, erlotinib), p<0.05 (0 compared to 2 μM erlotinib), p<0.0001 (0 compared to 50 nM, BGJ398), p<0.05 (0 compared to 1200 nM BGJ398), Tukey’s multiple comparison test. The double treatment for this cell line is shown in Figure 4E. (**B**) Cell viability (Cell titer Glo) of A549 cells grown in a range of trametinib (left graph) and BGJ398 (right graph) concentrations. Cell viability was normalized to DMSO control (n=3). Error bars represent s.d. p<0.02 (0 compared to 1200 nM BGJ398), Tukey’s multiple comparison test. The double treatment for this cell line is shown in Figure 4M. (**C**) Cilia length quantification of H23 cells treated with or without the FGFR inhibitor BGJ398 for 48 hours. Note that after treatment with BGJ398 cilia length was reduced. n = 150, error bars represent s.d. p<0.02, unpaired T test. (**D**) Cilia percentage quantification of H23 cells when treated with or without the FGFR inhibitor BGJ398 for 48 hours. Note that after treatment with BGJ398 cilia percentage was reduced. n = 150, error bars represent s.d. p<0.03, unpaired T test. (**E**) Cell viability (Cell titer Glo) of H23 grown in a range of trametinib concentrations. (**F**) Cell viability was normalized to DMSO control (n=3). Error bars represent s.d. Cell viability (Cell titer Glo) of H23 grown in a range of concentrations of the FGFR inhibitor BGJ398. Cell viability was normalized to DMSO control (n=3). Error bars represent s.d. p<0.02 (0 nM vs 300 nM), p<0.0001 (0 nM vs 900 nM), Tukey’s multiple comparison test. (**G**) Cell viability (Cell titer Glo) of H23 cells treated with trametinib (6.25 nM), BGJ398 (300 nM), or a combination of both. n = 3, cell viability is normalized to the DMSO control. Error bars represent s.d. p<0.0003, for an unpaired T test. (**H**) Cilia percentage quantification of H1792 cells treated with or without BGJ398 for 48 hours. Note that after treatment with BGJ398 cilia percentage was reduced. n = 150, error bars represent s.d. p<0.04, unpaired T test. (**I**) Cell viability (Cell titer Glo) of H1792 grown in a range of trametinib concentrations. Cell viability was normalized to DMSO control (n=3). Error bars represent s.d. (**J**) Cell viability (Cell titer Glo) of H1792 grown in a range of concentrations of BGJ398. Cell viability was normalized to DMSO control (n=3). Error bars represent s.d. p<0.0001, Tukey’s multiple comparison test. (**L**) Cell viability (Cell titer Glo) of H1792 cells treated with trametinib (6.25 nM), BGJ398 (300 nM), or a combination of both. n = 3, cell viability is normalized to the DMSO control. Error bars represent s.d. p<0.0001, for an unpaired T test. (**L**) Cell viability (Cell titer Glo) of H2228 parental cells treated with NVP TAE (0.5 μM), BGJ398 (1.2 μM), or a combination of both. Cell viability is normalized to the DMSO control. Error bars represent s.d. p<0.0001, for an unpaired T test. (**M**) Cell viability (Cell titer Glo) of H2228 NVP TAE resistant cells with NVP TAE (0.5 μM), BGJ398 (1.2 μM), or a combination of both. Cell viability is normalized to the DMSO control. Error bars represent s.d. p<0.002, for an unpaired T test.

**Supplementary figure 7.**
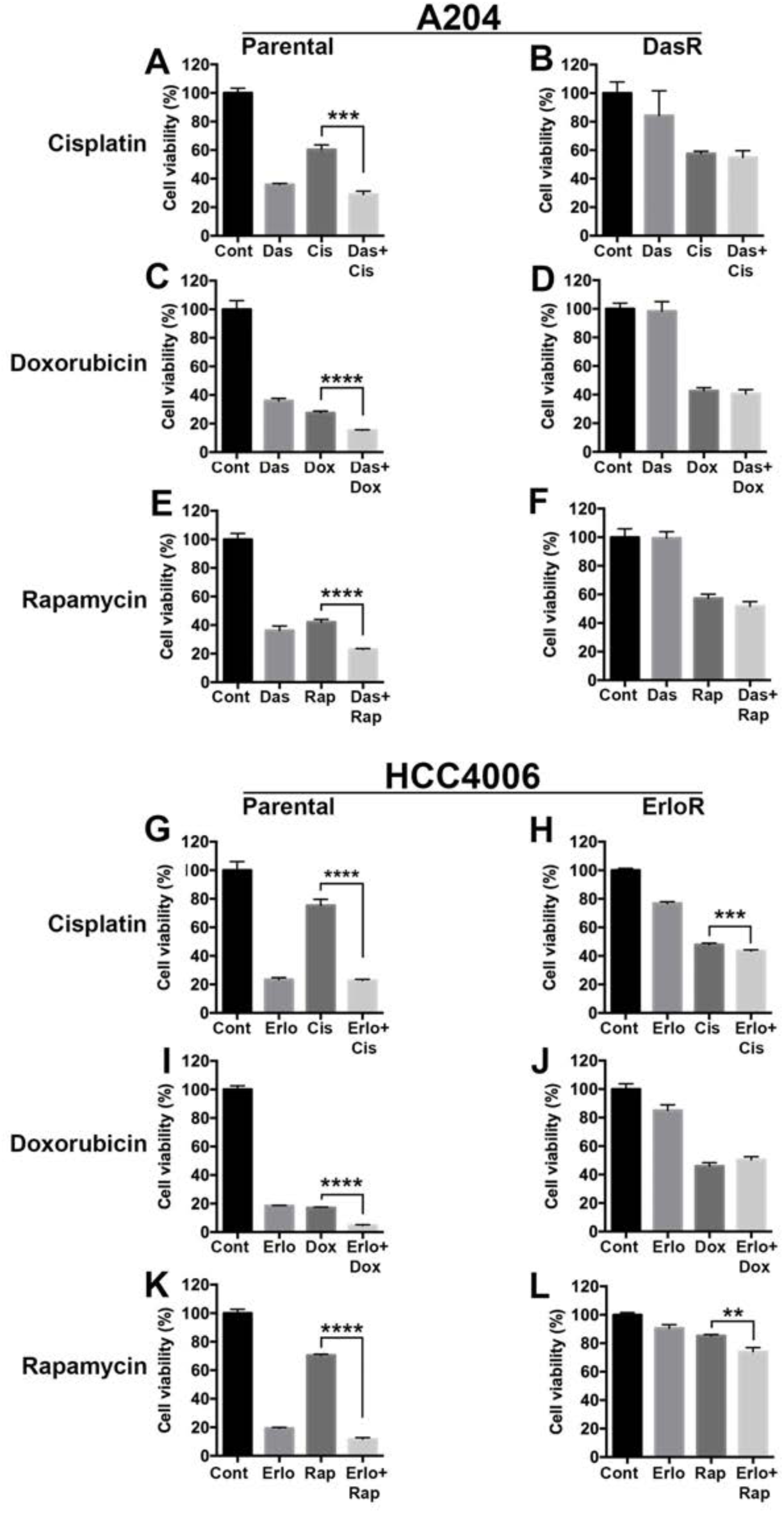
Cell cycle arrest does not sensitize HCC4006 and A204 kinase resistant cells to kinase inhibitors. (**A, B**) Cell viability (Cell titer Glo) of A204 (**A**) and DasR (**B**) cells treated with the S phase cell cycle inhibitor cisplatin (1 μM), dasatinib (0.5 μM), or a combination of both.n = 3, cell viability is normalized to the DMSO control. Error bars represent s.d. p<0.0002 (**A**), unpaired T test. (**C, D**) Cell viability (Cell titer Glo) of A204 (**C**) and DasR (**D**) cells treated with the G2/M phase cell cycle inhibitor doxorubicin (0.1 μM), dasatinib (0.5 μM), or a combination of both. n = 3, cell viability is normalized to the DMSO control. Error bars represent s.d. p<0.0001 (**C**), unpaired T test. (**E, F**) Cell viability (Cell titer Glo) of A204 (**E**) and DasR (**F**) cells treated with the G1 phase cell cycle inhibitor rapamycin (0.5 μM), dasatinib (0.5 μM), or a combination of both. n = 3, cell viability is normalized to the DMSO control. Error bars represent s.d. p<0.0001 (**E**), unpaired T test. (**G, H**) Cell viability (Cell titer Glo) of HCC40006 parental (**G**) and ErloR (**H**) cells treated with the S phase cell cycle inhibitor cisplatin (15 μM), erlotinib (0.5 μM), or a combination of both. n = 4, cell viability is normalized to the DMSO control. Error bars represent s.d. p<0.0002 (**H**) p<0.0006 (**H**), unpaired T test. (**I, J**) Cell viability (Cell titer Glo) of HCC40006 parental (**I**) and ErloR (**J**) cells treated with the G2/M phase cell cycle inhibitor doxorubicin (1 μM), erlotinib (0.5 μM), or a combination of both. n = 3, cell viability is normalized to the DMSO control. Error bars represent s.d. p<0.0001 (**I**), unpaired T test. (**K, L**) Cell viability (Cell titer Glo) of HCC40006 parental (**K**) and ErloR (**L**) cells treated with the G1 phase cell cycle inhibitor rapamycin (1 μM), erlotinib (0.5 μM), or a combination of both. n = 3, cell viability is normalized to the DMSO control. Error bars represent s.d. p<0.0001 (**K**), p<0.004 (**L**) unpaired T test.

**Supplementary figure 8.**
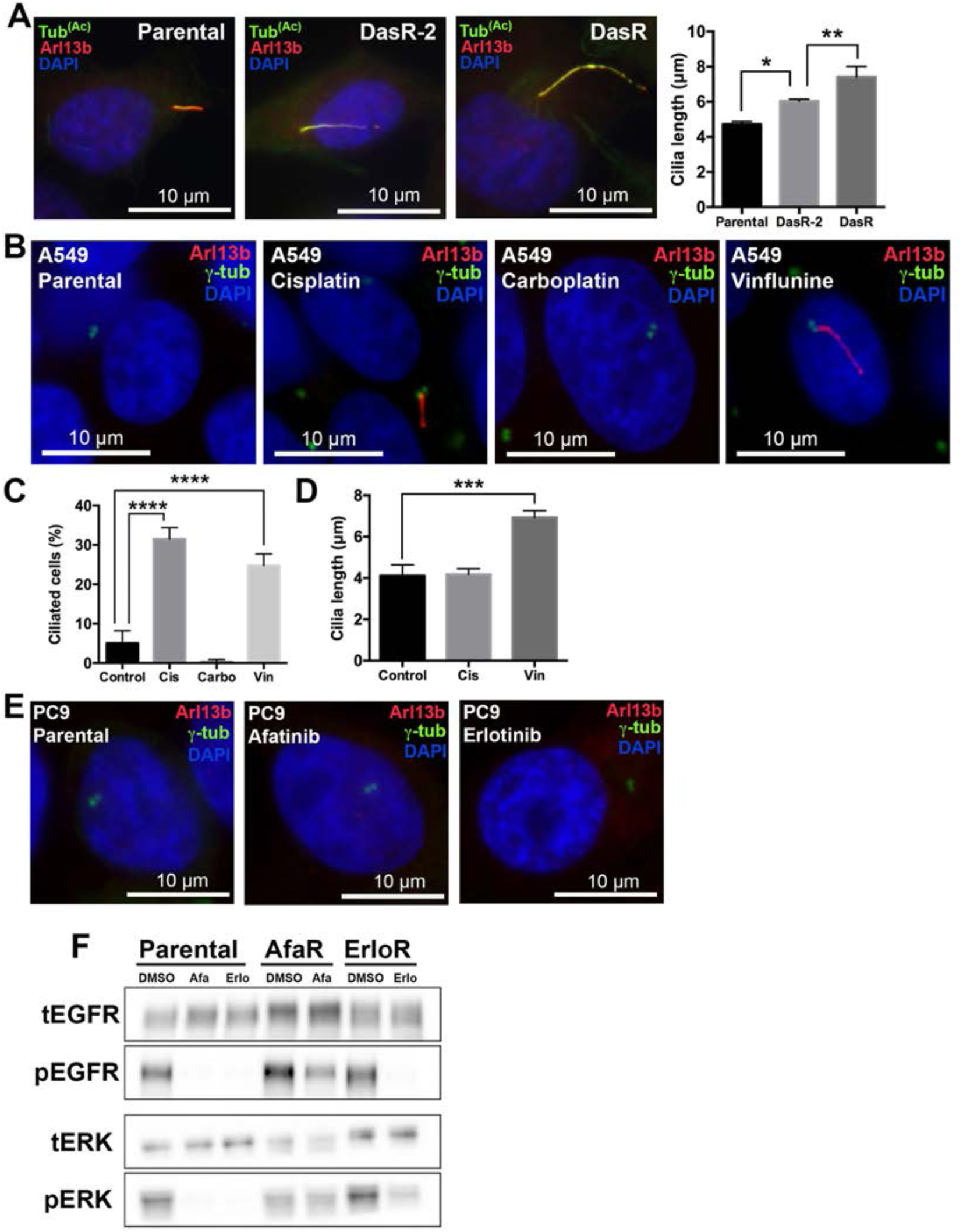
Ciliogenesis in additional models of acquired drug resistance. (**A**) Rhabdoid tumor A204 cells and two independently derived dasatinib resistant sublines (DasR and DasR-2) were stained with acetylated tubulin to mark cilia (green), Arl13b (red) and with DAPI (blue). Cilia length quantification is shown on the right. (n = 150), error bars represent the s.d. p<0.02 (parental vs DasR-2) and p<0.001 (DasR-2 vs DasR), Tukey’s multiple comparison test. (**B**) A549 parental cells and sublines resistant to cisplatin (cis), carboplatin (carbo) and vinflunine (vin) were stained with Arl13b to mark cilia (red),γ-tubulin (green) and with DAPI (blue). Note the increased ciliogenesis in sublines resistant to cisplatin and vinflunine. (**C, D**) Quantification of ciliated cells (**C**) and cilia length (**D**) shown in **B**. n = 300 (**C**), n = 150 (**D**). Error bars represent s.d. p<0.0001 (**C**), p<0.004 (**D**) Tukey’s multiple comparison test. (**E**) PC9 parental cells and sublines resistant to afatinib and erlotinib were stained with arl13b to mark cilia (red),γ-tubulin (green) and with DAPI (blue). (**F**) Western blot showing phosphorylated EGFR, total ERFR, phosphorylated ERK and total ERK in PC9 parental cells, afatinib (AfaR) and erlotinib (ErloR) resistant sublines. Cell were treated with or without afatinib (2 μM) or erlotinib (1 μM) for 3 hours when indicated.

**Supplementary Table 1.**
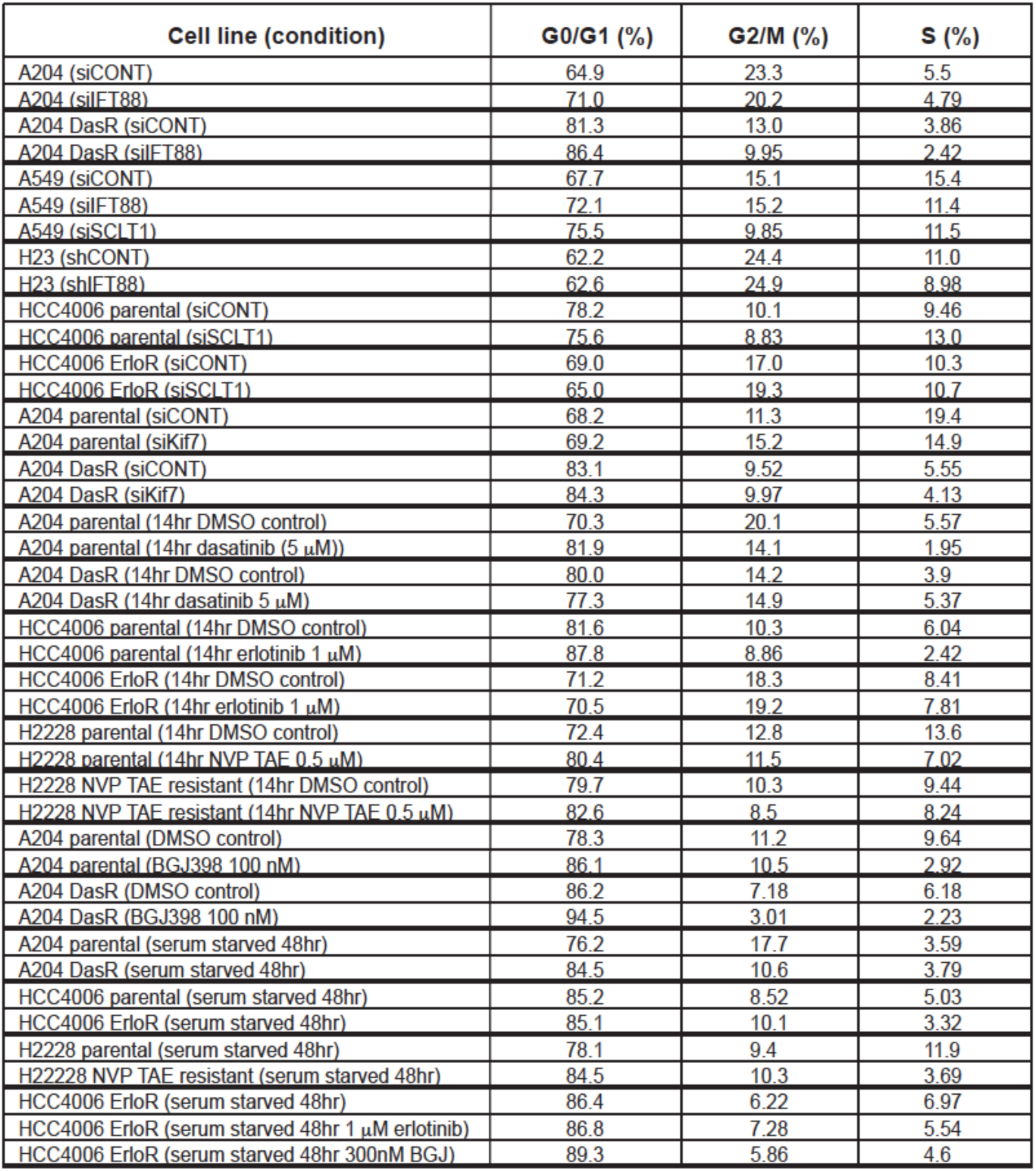
Cell cycle distributions for experiments shown. Note the minimal increase in cells in G0/G1 in A204 DasR, H23 and HCC4006 ErloR in response to siIFT88 compared siCONT. In addition siSCLT1 (A549) and siKif7 (A204) had minimal impact upon the cell cycle. Kinase inhibitors were tested in isogenic pairs, note the resistant sublines (HCC4006 ErloR, A204 DasR and H2228 NVP TAE resistant) had minimal changes to their cell cycle in response to their corresponding drugs (compared to DMSO controls).

